# PeakATail: precision-first poly(A)-site calling and calibrated alternative polyadenylation analysis in single-cell RNA-seq

**DOI:** 10.64898/2026.09.18.752791

**Authors:** Amir Amiri Tabat, Elyar Bafandeh Zendeh, Yasin Kaymaz

## Abstract

Most human genes can end their messenger RNAs at more than one position, and the choice shifts between cell types. Single-cell sequencing reads that choice in individual cells, but two questions are seldom asked: are the reported sites real, and does the test for a cell-type difference hold its error rate? Here we introduce our tool PeakATail, which calls polyadenylation sites from reads still carrying a piece of the tail, ranks them by how many distinct molecules carry that evidence, discards sites where adenosine-rich genomic sequence mimics a tail, and pairs them with a differential test whose error rate is measured, not assumed. The evidence is scarce, so the tool is built to be right rather than complete. Across four libraries from two species, 71 to 83 percent of its calls fall within 100 bases of a curated atlas, over thirty times the shuffled-control rate; long reads from different donors support 76.5 percent of the human calls. The cost is sensitivity. Only one of six test configurations held its false-positive rate under shuffled labels; with it, 15, 942 cell-type switches replicated across twelve lung cancer patients and none in ten shuffled-label runs. One pre-registered target was missed and is reported.

## Introduction

Most mammalian genes carry more than one polyadenylation site (PAS), and the choice among them, alternative polyadenylation (APA), rewrites the transcript’s 3’ untranslated region, changing its stability, localization and translational output (1, 2). The choice tracks cell state: proliferating and transformed cells favour proximal sites and shorter 3’UTRs (3, 4), and differentiation programs (spermatogenesis above all) remodel 3’-end usage wholesale (5). Droplet-based 3’ scRNA-seq primes on the poly(A) tail and sequences the transcript’s 3’ end, so every such library is, in principle, an untargeted single-cell 3’-end assay, and the raw material for placing PAS and testing their differential usage between cell types already sits in the public archives. Extracting it is harder than it looks. Oligo(dT) also primes internally at genomically templated A-rich stretches, producing spurious 3’ ends (6); and most reads stop short of the cleavage site, so a coverage pile-up marks a 3’-proximal neighbourhood, not a cleavage event.

A substantial tool ecosystem has grown around this opportunity; its members differ most fundamentally in their evidence model. Coverage-shape callers infer sites from the geometry of the read pile-up: Sierra fits splines to per-gene 3’ coverage to call peaks and couples them to a DEXSeq-based differential-usage test (7), and scAPAtrap combines coverage peak finding with poly(A)-tail-anchored reads to sharpen peaks toward nucleotide resolution (8). Read-evidence callers require a signature of cleavage in the reads themselves: polyApipe groups reads carrying non-templated poly(A) soft clips into poly(A)-site ranges (9); SCAPTURE calls peaks de novo and then evaluates them with the DeepPASS deep-learning sequence classifier (10); and scTail reads the cleavage position directly from the cDNA end of read 1, filtered by a pretrained sequence model, in library types that preserve a cDNA-bearing read 1 (11). Catalog-based methods sidestep de novo calling altogether: scUTRquant quantifies against a curated, cleavage-site-augmented transcriptome, gaining positional fidelity at the price of seeing only catalogued sites (12).

What the ecosystem still largely lacks is a way of knowing how much of a PAS call set, and of the ranked switch list computed from it, deserves belief. This paper attacks two reliability gaps. The first is the calibration of differential tests: tools ship differential-usage statistics, but their behaviour under a true null is rarely measured; UMI pseudoreplication, marker-gene double-dipping and fragile dispersion estimates can each make a nominal false-discovery rate optimistic, and the recipient of a q-value cannot see any of this from the output. The second is benchmark circularity. De novo callers are conventionally scored by agreement with curated atlases such as PolyASite 2.0 (13). That is a reasonable proxy, but one in which an atlas-novel true site counts as an error, and one that structurally rewards methods whose priors resemble the atlas; we show below that a scoring model tuned to the atlas can beat a simple rule on atlas agreement while losing against atlas-independent long-read truth. Evaluation choices compound the problem: shipped call-set sizes in the benchmark panel assembled here span 22-fold on a single PBMC library, so a single-operating-point comparison measures the call budget as much as the caller, and published benchmarks (14, 15) rarely control for this with matched call counts, shuffled-coordinate nulls or truth independent of the atlas.

PeakATail (command-line name: `peakatail`) is built so that both gaps become measurable. It seeds PAS from direct poly(A) evidence: reads whose non-templated poly(A) soft clips record cleavage and polyadenylation on the individual molecule. The channel is small but highly specific; in the deep 10x v3 PBMC library used here, only 0.573% of accepted cell-barcoded reads carry a qualifying poly(A) clip. PeakATail therefore emits tiered output, keeping molecule-counted, clip-supported sites separate from coverage-only summits (which are never switch-tested in this paper), and applies a strand-aware internal-priming filter. Around the caller we built the evaluation the field’s tools have not had: acceptance gates and definitions registered before the final run existed; gene-body-shuffled nulls beneath every headline metric; an atlas-independent check against poly(A)-verified long-read 3’ ends (PacBio Kinnex; 16) from donors other than the short-read library, stated as such; a switch test whose false-discovery-rate control is measured under label permutation rather than assumed; and a replication filter that reports a cell-type switch only when it recurs, in the same direction, across independent patients, with the entire pipeline re-run under patient-wise label shuffles.

Pre-registration is informative only if failure is a reportable outcome, and this paper’s negative results are part of its contribution: a pre-registered "trusted novel PAS" definition that missed its long-read acceptance target and is reported as a failure; a clustering-based novelty claim withdrawn after its own ablation; a literature-derived gene panel that does not reproduce; and our own shipped differential-test defaults, which do not control the false-discovery rate. A pre-registration that can only be confirmed is decoration, and a literature in which no single-cell APA caller has ever reported a failed acceptance gate is a literature in which acceptance gates are not being set.

The claim of this paper, in one sentence: PeakATail calls poly(A) sites from direct poly(A) evidence with the highest atlas-agreement precision of any de novo tool benchmarked, couples them to an FDR-calibrated switch test, and demonstrates (under pre-registered gates, a shuffled-null discipline and an atlas-independent long-read check) that its cell-type APA switches replicate across patients, with zero replication in label-shuffled nulls.

## Materials and methods

### Datasets and reference resources

No new sequencing data were generated. Human PBMC, donor 1: the 10x Genomics public library `pbmc_10k_v3` (Single Cell 3′ v3 chemistry; CellRanger 3.0.0 (17) on the GRCh38-3.0.0 reference bundle; 91 bp read 2; ∼10, 000 estimated cells; 557, 564, 408 accepted cell-barcoded reads). Human PBMC, donor 2: the 10x public library `pbmc4k` (Single Cell 3′ v2; CellRanger 2.1.0 on GRCh38-1.2.0; 98 bp read 2; 4, 340 estimated cells; mean 87, 433 reads per cell; flow cell H53GNBCXY). 10x publishes no donor identifier for `pbmc4k`: it is demonstrably a second library, chemistry, flow cell and CellRanger version, and only presumptively a second individual. Mouse testis: the two adult spermatogenesis libraries of GEO GSE104556 (21, 22; Single Cell 3′ v2; 98 bp read 2; 1, 294 and 1, 364 stage-labelled cells), aligned with STARsolo (18, 19). Cohort: the Laughney lung-adenocarcinoma atlas, GEO GSE123904 (23, 24), comprising 17 libraries from 14 patients (three patients contributed a tumour and a normal library; ∼224 GB of BAM); 29, 063 curated cells entered the switch pipeline, 18, 651 (64.2%) surviving label confirmation (below). Annotation builds are Ensembl 99 (GRCh38) and Ensembl 102 (GRCm38); the accuracy reference is the PolyASite 2.0 representative-site atlas (13; 569, 005 sites on GRCh38, 301, 006 on GRCm38). Per-library parameters, pipelines and poly(A) clip rates are in Fig S1; unmeasured values are shown as missing, never approximated.

### The caller and the pre-registered default output

The formal statement of every definition in this section, in notation, is the formal specification in the Supplementary Data; where the two are read together, this section governs scope and that file governs definition. PeakATail was run at two frozen commits: v1 `4efeb125` and v2 `9dfdefb3eb353b0817ef79c4eb9ace6d6c8aab53`. Each was run from a dedicated worktree whose path every run manifest asserts (test suite at v2: 1, 277 passed). All current numbers come from the v2 runs; v1 values are labelled comparisons.

### Read acceptance

A read is used when it is mapped, on the strand of the current pass, carries a 16-nt cell barcode tag and aligns over at most `--seq-len` nucleotides (91 bp for `pbmc_10k_v3`, 98 bp for `pbmc4k` and both mice, set to the library’s read-2 length). The length test compares the alignment’s *reference* span with `--seq-len`, so spliced alignments whose genomic span exceeds the read length are excluded before clip detection and counting: 13.74% of valid-cell-barcode reads genome-wide on `pbmc_10k_v3`, 96% of them spliced. This is a documented, conservative limitation: relaxing it to a query-length test adds only +1.76% clip reads (projected ∼+0.4% recall) and is roadmap work that ships in no result here.

### Clip detection

A read enters the poly(A)-evidence channel when its 3′-side terminal soft clip is ≥6 nt (`--polya-min-clip 6`) and ≥80% A on + / T on `−` (`--polya-min-purity 0.8`), with a ≥6-nt run flush to the alignment edge. By these criteria 0.573% of accepted cell-barcoded reads in `pbmc_10k_v3` qualify genome-wide (3, 195, 067 / 557, 564, 408). An earlier figure of 1.152% is corrected: it was an artefact of a QC estimator that samples only the first 200, 000 cell-barcoded reads of a coordinate-sorted BAM, the head of chr1, and appears here only as a superseded estimate.

### Coverage candidate regions

Clip clusters are seeded against a coverage pass over the same reads, which supplies the tier-2 candidates and the count source for tier-1 sites. Accepted read 3′ ends are accumulated per (contig, strand); a candidate region opens where at least `--default-threshold` (5) ends overlap and closes when an incoming read’s start passes the position of the fifth-most-recent end, and a region starting within `--merge-len` (100 bp) of the previous one is joined to it. `--dynamic-threshold` is off in every run reported here, so that opening threshold is the fixed 5 rather than a local-density estimate. Within a region the production coverage strategy — `lambda_gradient`, the strategy that `clip_seeded` wraps by default — gates the region twice before localising anything: its maximum height must be ≥20 reads, and it must clear a Poisson test at α = 0.05 against a local background λ taken as the median of the positions below 10% of the regional maximum. A region that fails either gate yields no site at all. Surviving regions are smoothed with a 50 bp centred moving average, and the indices where the first difference turns from positive to non-positive are the candidate summits. A summit is kept when its raw height is ≥20 reads, strictly above λ, individually Poisson-significant at the same α, and topographically prominent by ≥5 coverage units. Kept summits are ranked by prominence × height/λ plus the smoothed second difference, the 5 highest are retained, and the region is partitioned at the midpoints between them so every read end belongs to exactly one candidate. Two fallbacks are part of the definition: if the smoothed profile has no local maximum, or if no summit survives the filters, the whole region is returned as a single candidate. Adjacent candidates are then merged when their summits lie closer together than the library’s median read length (auto-detected, `--min-pas-spacing -1`) or when the valley between them falls below λ, which for the λ-based strategies supersedes the user-facing `--min-pas-prominence`. One scope note travels with these values: the coverage strategy is instantiated at its own defaults, so the `--smoothing-window`, `--min-prominence` and `--max-pas` values recorded in each run configuration are not forwarded into it; in every run reported here those recorded values equal the defaults given above (50, 5.0, 5), so no number in this paper differs either way. The candidate count this stage delivers on `pbmc_10k_v3` is the entry point of the funnel in Fig 1.

**Fig 1.**
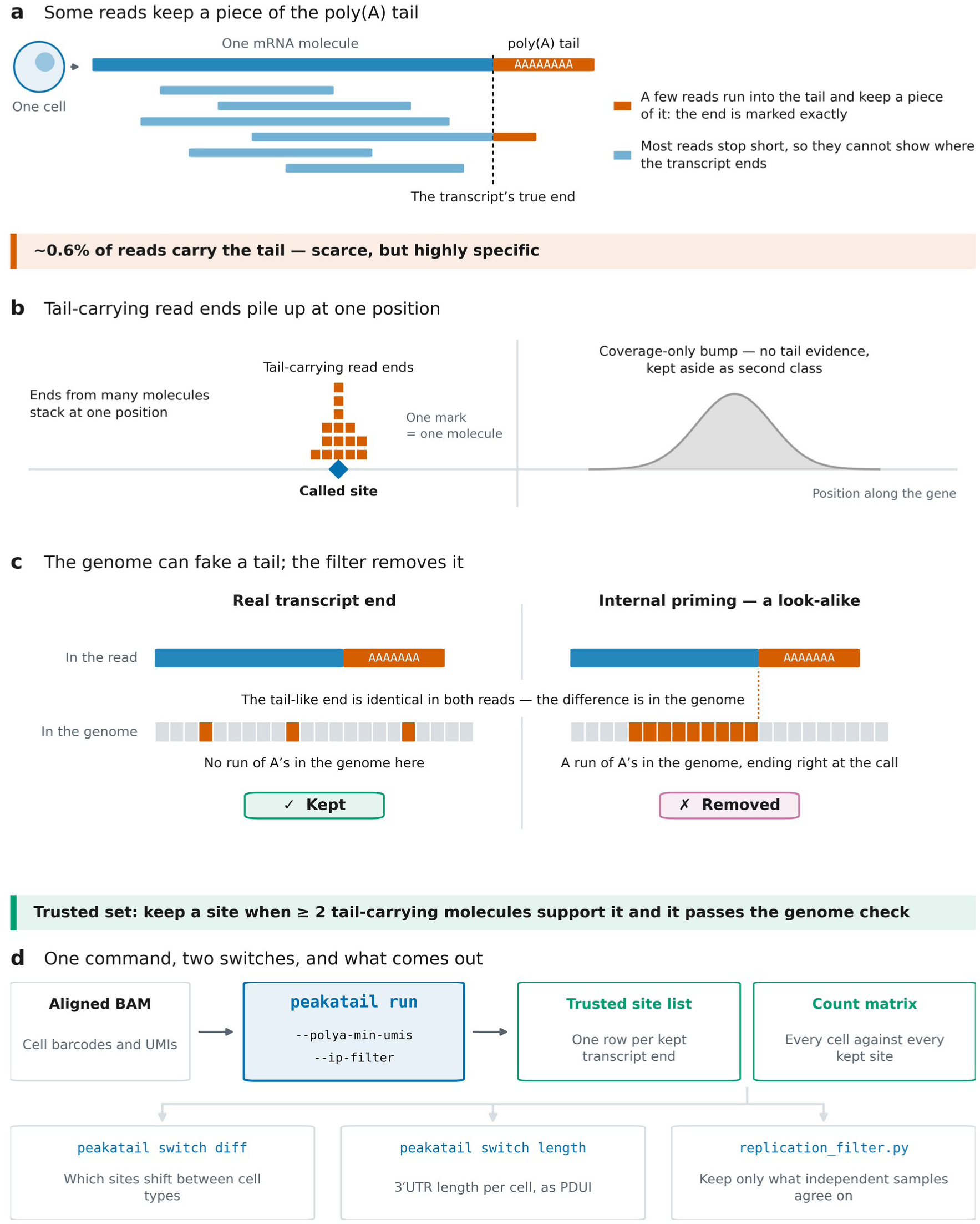
How PeakATail finds the 3′ ends of transcripts in single cells, and what it gives you. All four panels are schematics drawn to explain the method: no panel plots a data series, and every quantity a panel stands for is measured by the figure’s script and written to its audit table (code 9dfdefb; PBMC 10k v3 arm). **(a)** An mRNA ends in a poly(A) tail that is added after cleavage and is not written in the genome. Most reads stop short of that end, but a small fraction run into the tail and keep a piece of it as a soft-clipped A-run; because that piece is not genomic, the read’s last aligned base is the transcript’s end at single-base resolution. Genome-wide, 0.5730% of accepted cell-barcoded reads carry such a piece (3,195,067 of 557,564,408). **(b)** The last aligned bases of tail-carrying reads from many different molecules stack at one position; PeakATail joins them into a cluster and calls its read-weighted mode a site, with support counted in distinct (cell barcode, UMI) molecules. A coverage summit that no tail-carrying read claims is written to a separate, second-class file and is not in the default output: its atlas agreement is 0.0558 against 0.7062 for the default. **(c)** An A-rich stretch encoded by the genome is copied into the RNA and can capture the oligo(dT) primer, giving a read whose clipped A-run is indistinguishable from a tail-carrying read’s (internal priming). The A-rich genome lies upstream of and at the called base, not downstream of it: the aligner runs into the tract and clips where it ends, so the run ends at or before the call in 91.7% of removals. The genome tells the two apart, so PeakATail reads the 40 nt of genome sequence around every called end (offsets −9…+30) and removes calls that sit on a run of genomic A. On PBMC 10k v3 that removes 15,588 of 62,132 ≥2-molecule calls and moves atlas agreement from 0.5907 to 0.7062; it also costs recall, since 24.6% of what it removes does have an atlas site within 100 bp. **(d)** The tool. One command, `peakatail run`, takes an aligned single-cell BAM carrying cell barcodes and UMIs and returns the trusted site list and a matrix of every cell against every kept site. Two switches set the operating point: `--polya-min-umis`, the number of distinct tail-carrying molecules a site must have, and `--ip-filter`, the genome check of panel c. Neither switch is drawn with a value, because `--polya-min-umis` resolves to 1 at the caller while the badge above states the paper’s pre-registered ≥ 2-molecule OUTPUT rule; printing the caller’s 1 beneath that badge would read as a contradiction. Three commands then read the same matrix: `peakatail switch diff` for sites that shift between cell types, `peakatail switch length` for 3′UTR length per cell as PDUI, and `replication_filter.py` to keep only what independent samples agree on. The panel is a schematic of the interface and deliberately plots no measured quantity: the accuracy of these outputs is Figs 2 and 3, the false-positive rate of `peakatail switch diff` (3.0% of label-shuffled tests p < 0.05, 0/20 null runs with a q < 0.05 hit) is Fig 4. Caveats that travel: atlas agreement is agreement with a curated atlas, not ground truth, so an atlas-novel true site counts as a false positive, and the recall denominator is the atlas restricted to genes detected in the dataset; the clip rate is one PBMC donor and one chemistry (CellRanger 3.0.0, 10x 3′ v3, 91 bp read 2), and any pipeline that trims poly(A) before alignment destroys this evidence channel entirely; the tier label reflects clip support, not gene assignment (PAS-to-gene assignment in overlapping loci was defective in the code behind this figure, repaired upstream only after 9dfdefb and with a residual acknowledged there, so no gene-level claim is made here); neither rule the figure names is a shipped command-line default, since `--polya-min-umis` is 1 at the caller and ≥2 molecules is the pre-registered *output*, while the genome check is the opt- in `--ip-filter`, run here in filtering mode, so the figure labels this the trusted set and not the default; and the four panels are schematics that must not be read as data: the evidence for every claim they make is in Figs 2 and 3 (accuracy, resolution and reproducibility), Fig 4 (calibration) and the audit tables. Alt text: Four-panel schematic diagram, no plotted data. Panel a, a messenger RNA whose genome-encoded body ends at a cleavage point followed by a non-templated poly(A) tail, with a sequencing read running into the tail and retaining a soft-clipped run of adenosines. Panel b, the last aligned bases of many such reads from different molecules stacking at one genomic position, which is called as a site. Panel c, an adenosine-rich stretch written in the genome capturing the oligo(dT) primer and producing a look-alike clipped read, with the adenosine run lying at or upstream of the called base rather than downstream. Panel d, a flow diagram of the software: one calling command takes an aligned single-cell BAM and returns a site list and a cell-by-site matrix, which three downstream commands then read.

### Clustering and tiers

Qualifying clip positions are clustered per (contig, strand) by single linkage with a 25 bp maximum gap between consecutive positions; the reported cleavage position is the read-weighted mode of the cluster. Clip-supported clusters form tier 1; coverage summits claimed by no clip cluster form tier 2, written to a separate low-confidence file (BED score 0). Support (BED column 5) is counted as distinct (cell barcode, UMI) molecules. Quantification additionally credits a tier-1 site with accepted read ends in its cleavage window ([site − seq-len, site + 25]) and with the counts of the coverage candidates it suppresses, so the count column is not the clip evidence. Cell filtering (`--min-read 1500`, `--min-cells 3`, `--min-pas-per-cell 50`) precedes matrix export, and `pasbed.bed` omits gene-assigned sites left with zero counts after it (0.45% on `pbmc_10k_v3`, 3.7% on mouse 1); the scored call sets are the post-cell-filter sets.

### Internal-priming (IP) filter

For each called site the genomic sequence in a transcript-relative window of −9..+30 nt (40 nt, anchored on the BED `end` for + and the BED `start` for `−` strand) is tested; a site is flagged when it contains ≥6 consecutive genomic A or an A-fraction ≥0.70 ( `--ip-a-stretch 6`, `--ip-a-fraction 0.7`). At v2 the veto itself was opt-in behind `--ip-filter`, and `--ip-filter-mode` defaulted to `annotate`, which keeps every site and only records the flag; every benchmark arm therefore passed `--ip-filter --ip-filter-mode filter` explicitly, which removes flagged sites. Both defaults have since changed on `develop`, after every run reported here: `--ip-filter-default auto` runs the veto whenever a `--genome-fasta` is supplied, and `--ip-filter-mode auto` resolves to `filter`. That is the breaking change behind the 0.2.0 to 0.3.0 version bump. Because every arm here names both flags explicitly, the call sets scored in this paper are identical under either default, and no number reported here was produced with the new one. A correctness fix in v2 corrected the minus-strand anchoring: the forward-coordinate window had been applied to both strands, testing minus-strand sites over the wrong sequence. The fix leaves plus-strand output byte-identical and, on `pbmc_10k_v3`, newly removes 5, 599 and newly keeps 11, 217 minus-strand sites. The same v2 change replaced dense TF-IDF (the Signac method; 25) with sparse operations, parallelised peak calling per (contig, strand) with a deterministic merge, and streamed the cell-barcode filter, verified byte-identical on 18/18 compared outputs.

The pre-registered default output is tier 1 ∩ IP-pass ∩ ≥2 distinct clip molecules, filtered from the tier-1 file (`--polya-min-umis` remains 1 at the caller; ≥2 molecules is the pre-registered *output*). The ≥2-molecule threshold is a reliability choice, not the F1 optimum on any dataset tested; the ≥1-molecule tier-1 output is reported alongside as the sensitivity arm.

### Benchmark protocol

All accuracy numbers come from one scorer, `score_tool.py` (analysis repository), applied identically to every tool. Call sets are reduced to strand-aware 1-bp points (the 3′-most base: `end − 1` on +, `start` on `−`) and matched to PolyASite 2.0 representative sites by strand-matched point distance (`bedtools closest -s -d -t first`; 26; `LC_ALL=C` throughout). Atlas-agreement precision at window W (P@W) is the fraction of scored calls with a strand-matched atlas site within W bp (W = 10, 25, 50, 100, 200); it measures agreement with a curated atlas, not ground truth: atlas-novel true sites count as false positives. Detected-gene recall (R_det@100) is the fraction of the atlas restricted to genes detected in the dataset that is recovered within 100 bp; the denominators are 285, 136 sites (14, 949 genes, `pbmc_10k_v3`), 126, 686 (GSE104556) and 268, 097 (13, 763 genes, `pbmc4k`). Each denominator is built once per dataset by one recorded recipe and then shared, fixed, by every tool: detected gene IDs are read from a single PeakATail run’s `annotatedpas.bed` on that dataset, in which each site takes the nearest same-strand annotated gene by signed distance (zero inside the gene body, ties to the first gene in coordinate order) and is kept when it lies within that gene’s annotated 3′UTR length or within `--utr-multiplier` (2.0) times it, sites further out to `--max-gene-distance` (5, 000 bp) being the extended class that `--include-extended` leaves off in every run here (for `pbmc_10k_v3`, the earlier coverage-strategy `lambda_gradient` run, not the benchmarked clip-seeded arm; for `pbmc4k`, its only run, a declared deviation), gene bodies are restricted to those IDs, and the atlas is intersected strand-matched. Because the detection source is a PeakATail run, the construction’s sensitivity to that choice was measured on PBMC, where two runs exist: rebuilding from the clip-seeded v2 run gives 15, 271 genes / 288, 740 sites versus the shipped 14, 949 / 285, 136 (+1.26% denominator size, 99.5% of shipped sites in common), and a larger denominator lowers R_det, so the choice biases against the arm supplying the detection; the donor-2 recall gap additionally survives scoring both arms against both donors’ denominators. Full-atlas recall (569, 005 / 301, 006 sites), which involves no detection choice, is reported next to every R_det; F1_det is the harmonic mean. Each call set is compared against its own null: 3 seeds of width-preserving `bedtools shuffle -chrom -noOverlapping` inside merged detected-gene bodies. Scored n sits slightly below raw BED line counts because the scorer drops non-primary contigs. The v2 gate table was reproduced to six decimals by an independent `bedtools`-based reimplementation.

### Competitors

Sierra (7), scAPAtrap (8), polyApipe (9), SCAPTURE (10) and scUTRquant (12) were each run on the same BAMs on the same machine under their own documented workflows, and their call sets were scored exactly as above (versions, parameters, run logs and retry/skip caveats: Supplementary Table T7 and Fig S6). scUTRquant is catalog-based and is shown but not ranked against de novo callers. scTail (11) could not be run on these libraries (read 1 is 28 bp). SCAPTURE’s mouse-2 site-level run completed and was scored (P@100 0.672); only its per-cell quantification step failed, which site-level benchmarking does not use.

### Matched-call-count comparisons

Because atlas-agreement precision and recall both move with call budget, no single-point head-to-head is quoted without both tools’ call counts. Each tool’s call set was truncated to a common N under its own shipped ranking (clip molecules for PeakATail, `peakdepth` for polyApipe, and Sierra’s and scAPAtrap’s own shipped per-peak rankings); within-tie ordering was re-scored under both orderings (coordinate order and ascending depth) on PeakATail’s own truncation, at four `pbmc_10k_v3` budgets (35, 759, 40, 519, 106, 170, 120, 916) and three mouse-1 budgets (22, 550, 23, 645, 35, 732), and the precision and recall leads over polyApipe survive both orderings at every one of them. Competitor-side ties were not re-ordered, so matched-N precision claims in the interval 20, 320 ≤ N ≤ 35, 759, where the competitor truncations cut inside tied groups, are reported as unestablished rather than as a lead. Two exclusions are principled: SCAPTURE ships no non-circular ranking column, so it cannot be truncated; and scAPAtrap truncations below its 19, 410-call score cap cut arbitrarily inside a tied group and are reported as such, never as a real ranking.

### Long-read truth construction

The atlas-independent read-out uses publicly available PacBio Kinnex (16) long-read PBMC data from the x3p and GEM-X library preparations, distributed by Pacific Biosciences as the public datasets `DATA-Revio-Kinnex-PBMC-10x3p` (Revio, 10x 3′ v3.1) and `DATA-Revio-Kinnex-PBMC-10kcells-10xGEMX3p` (10x 3′ v4 GEM-X) at https://downloads.pacbcloud.com/public/dataset/Kinnex-single-cell-RNA/ (downloaded 2026-08-19; files `scisoseq.5p--3p.tagged.refined.corrected.sorted.dedup.bam` and `scisoseq.mapped.bam` respectively). These are vendor-distributed datasets with no repository accession and no DOI, so the derived truth and decoy point sets built from them are deposited in the companion repository’s archived snapshot, which is the citable object for this truth set. The long-read data come from a different donor from the short-read libraries, so this is site-level, donor-mismatched truth. Poly(A)-verified read 3′ ends were collapsed to strand-aware point BEDs at support thresholds of ≥5, ≥20, ≥100 and ≥500 records; support is an alignment-record count at the terminus (∼10.5% of records repeat a (cell barcode, UMI) pair there and were not de-duplicated), so the thresholds are slightly optimistic. A decoy set of termini flagged by Kinnex’s own internal-priming rule (A-rich +1..+18 window) provides the false-positive read-out. Concordance is the fraction of calls with a same-strand truth terminus within 25 bp, read against a gene-body-shuffled positional null (10 seeds); the build script and truth BEDs are in the analysis repository.

### Pre-registration practice

Every acceptance gate was committed before the run it judges, with git-verified commit times where a commit exists and the document’s own recorded date otherwise. The original two-sided benchmark gate (P@100 ≥ 0.38 and F1_det > 0.261) predates the Stage-2 run that failed it; the precision-first default and its added gate (P@100 ≥ 0.50 on `pbmc_10k_v3` and both mice) were committed to the analysis record (`manuscript/13_reliability_positioning.md`) at 01:18:59 on 2026-08-21, 1 h 41 m before the final arms started (02:59:36), and were unchanged for the v2 re-run (16:14). The ≥2-molecule threshold was informed by a post-hoc sweep of a superseded run before being pre-registered: a disclosed deviation, cited nowhere as a result. Later tests were pre-registered the same way: the trusted-novel definition and its 70% target, the second-library test, the `peakAtail-prime` adoption criterion, and a 3′UTR-singleton promotion rule (still untested; shipped flag-off). Amendments are appended, never rewritten; Amendment 3 records that the prime criterion was drafted against a v2 default configuration that never existed and that prime’s default output is byte-identical to the v2 manuscript arm (every pre-registered delta exactly 0.000000). Every prime-branch number is exploratory with respect to the manuscript’s gates.

### Differential APA test and its calibration

Cell-type APA switches are tested with `peakatail switch diff`: for each (cell-type pair, PAS), a Fisher exact test of PAS counts against the within-gene total across the two cell types, with `--count-mode cells` (a cell contributes at most once per PAS; `reads` mode counts UMIs and pseudoreplicates within cells) and marker pre-selection off (`--marker-top-n 0`), at the default aggregation scope `--isoform-agg per_gene`, in which each PAS is tested against the remainder of its gene rather than against the other sites of its own 3′UTR isoform; Benjamini–Hochberg correction (20) is applied per cell-type pair within each library, so that each (library, pair) is one BH family, and |Δproportion| is reported as the effect size. Genes with a single tested PAS yield no table and are dropped.

Fisher on cells treats cells within a library as independent; ambient RNA, doublets and clonal structure violate that assumption, and the label-permutation null measures exchangeability under the global null only, so it cannot detect miscalibration arising from within-library dependence. The design’s defence is structural rather than distributional: no within-library switch is reported on its own, because the unit of replication is the patient, so within-library dependence cannot by itself manufacture a cross-patient replicated call; and the label-shuffle null pipeline preserves each library’s cells, counts and dependence structure while destroying the label association, then passes through the identical replication filter.

The configuration was chosen by a calibration harness: on testis mouse 1 (1, 230 cells with GEX-derived stage labels from marker-panel argmax on the STARsolo gene matrix, orthogonal to the PAS matrix under test; SPC 370 / RS 504 / ES 356; 76, 711 PAS; three stage pairs), six configurations (Fisher reads/cells × marker pre-selection on/off, and `nb_pairwise`) were each run on the true labels and on 20 label permutations shared identically across arms (6 × 21 runs, 0 failures). The pre-stated calibration rule (null p < 0.05 ≤ 7%, null p < 0.01 ≤ 1.5%, null q < 0.05 ≤ 5%, and ≤25% of null runs carrying any q < 0.05 hit) is our operational rule, committed in the figure script; Kolmogorov– Smirnov tests against uniformity are reported, not gated. Wherever marker pre-selection or `nb_pairwise` is used, permutation-calibrated q-values (empirical p ranked within pooled same-pair null p-values, BH per pair) are required. The calibrated configuration was re-validated on the final caller on both mice.

### Cohort switch replication (Laughney)

All 17 libraries were peak-called in one cohort-mode run ( `ema run -c` at the v2 commit, `peakatail run -c` from 0.3.0, `clip_seeded` with `--ip-filter --ip-filter-mode filter`, `--threads 16 --peak-workers 16`), so every library shares the tool’s own unified PAS identifier space (505, 197 sites), with no post-hoc coordinate harmonisation. The pre-registered tested universe (`tier1_ge2`) is IP-pass ∧ tier 1 in ≥1 member library ∧ cohort clip-molecule sum ≥2, giving 80, 464 PAS (40, 811 +, 39, 653 `−`); restricting to it changes the within-gene Fisher denominator (cells expressing any universe PAS of the gene) and drops genes with fewer than two universe PAS. Tier-2 sites are never switch-tested.

Cells enter the test only under the pre-registered label-confirmation policy: curated immune labels must agree with a CellTypist call (27; confidence ≥0.5, mapping table frozen before use); the four non-immune types (epithelial/tumour, fibroblast, endothelial, pericyte), which the immune model cannot confirm, require a canonical-marker gate on the cell’s own gene expression (EPCAM/KRT8/KRT18; COL1A2/DCN/LUM; PECAM1/VWF/CLDN5; RGS5/ACTA2/PDGFRB).

Unconfirmed cells are dropped, not relabelled; cell types with <20 confirmed cells in a library are dropped for that library. A cell-type pair is tested within each library where both types survive that floor; pooled across libraries the cohort tests 59 distinct cell-type pairs, of which 12 occur in only one patient and can never replicate (the replicated set spans the other 47). The switch test runs per library; ten patient-wise label-shuffle null runs per library go through the identical pipeline.

The unit of replication is the patient, not the library (either library of a two-library patient counts once toward that patient’s support; audited on the Macrophage–T cell pair, where collapsing to patients removes 89 PAS that would have "replicated" on two libraries of single patients). The pre-registered replication filter (`replication_filter.py` v0.2.0, md5-recorded) reports a (pair, PAS) switch only if called at q < 0.05 in the same direction in ≥2 patients, with any opposite-direction call vetoing it, including between the two libraries of one patient (audited: 2 vetoed PAS where one patient’s two libraries called opposite directions); |Δproportion| ≥ 0.1 is reported alongside as the effect floor, and a ≥3-patient sensitivity configuration is reported. The replication primary is 15 libraries from 12 patients: GSM3516664 (bone metastasis) is excluded by a pre-declared data-quality rule whose stated premise later proved to be a clip-rate sampling artefact of the head-of-BAM estimator, so the exclusion stands only as pre-registered, with a sensitivity run including it reported; and GSM3516671 retains one confirmed cell type, so it yields no testable pair. Null combinations are index-aligned; ten permutations floor the empirical p at 1/11 = 0.091, so the null yield is a null control ("none in 10 label-shuffle nulls, empirical p ≤ 0.091"), never an FDR estimate. Replication statistics are computed on PAS identifiers; no ranked list of named switch genes is reported, because the PAS-to-gene assignment in the frozen v2 code mis-assigns PAS in overlapping loci (repaired after the freeze, the repair carrying a documented residual tie-break case, and that fix is not in the code that produced these results).

### Spermatogenesis analysis

The v2 record reuses the frozen v2 benchmark runs of both mice (h5ad checksums verified against the run manifests; no caller re-run) with byte-identical GEX-derived stage labels (marker-panel argmax over Leiden clusters of the STARsolo gene-expression matrix, orthogonal to the PAS matrix under test; panel and assignment script in the companion repository; SPC/RS/ES = 370/504/356 and 268/757/271 cells; spermatogonia excluded as fragile). The per-gene statistic is PDUI = distal / (proximal + distal), where the proximal and distal member of each gene’s PAS pair are its first and last PAS in transcription order (strand from `pasbed.bed`, re-asserted per row), computed as the depth-weighted pseudobulk PDUI per stage. Per-gene 3′UTR usage across the SPC, RS and ES stages was computed on depth-guarded genes (≥50 UMI at the PAS pair in every stage; 1, 548 / 1, 200 of ∼10, 500 genes with a pair); the monotone-shortening fraction is read against 20 stage-label shuffles (z-score against the shuffle distribution), monotone lengthening is tabulated identically, and the direction-specific evidence is the shortening-minus-lengthening excess (binomial test). A composition-controlled per-cell distal-usage residual is compared between stages by Cliff’s δ against the full label-shuffle null range; per-gene medians and the across-gene UMI-weighted index are reported as descriptive scope checks. The calibrated switch test (five label shuffles per mouse) ran on all three stage pairs; cross-mouse replication matches q < 0.05 PAS between mice by gene and strand within 100 bp, requires the same direction, and is read against all 15 null pairings of label-shuffled runs, with per-gene effect sizes correlated across mice (Spearman ρ).

### Second-library generalisation test

The `pbmc4k` run was pre-registered before any number existed: the frozen v2 tree, the flag set copied verbatim from the `pbmc_10k_v3` arm, the same P@100 ≥ 0.50 gate. Of the 108 recorded parameters in `run_config.json`, 107 are identical between the two human runs; the only change is the library descriptor `--seq-len`, which moves from 91 to 98 (`--threads` differed, verified output-irrelevant), and nothing was retuned after seeing a `pbmc4k` number. Scoring was identical, with the library’s own detected-gene denominator built by the recorded recipe. Cross-library site concordance (each library’s default calls against a strand-matched default call of the other within 25 and 100 bp, both directions, with genic-shuffle nulls) was declared before computation; a matched-call-count control accompanies any cross-library atlas-agreement-precision comparison.

### Compute environment

All runs used one shared Linux server under `LC_ALL=C` (machine locale tr_TR), with wall time and peak resident set size (RSS) from `/usr/bin/time -v`; frozen worktrees and per-run manifests record the code commit of every run. Peak RSS is the production number, and wall times are quoted with their concurrency disclosed. On the `pbmc_10k_v3` BAM the v2 caller used 12.53 GB peak RSS at 34:37 wall (no-IP arm) and 11.12 GB at 32:59 (IP arm), measured with all four benchmark arms running concurrently (27:43 / 12.45 GB uncontended); the mouse arms used 3.07/3.65 GB at 14:10/16:00 (concurrent); the v1 PBMC figures were 293.7 GB / 3:45:53 before the memory fix. The 17-library cohort run took 1:06:26 at 13.53 GB on v2 (9:06:26 / 13.63 GB on v1) at identical output scale (505, 197 unified PAS on both codes). The `pbmc4k` run (8 threads, 15:41, 3.24 GB) is a run record, not a benchmark. Competitor wall times are their own runs on the same machine and carry their retry/skip caveats verbatim (Fig S6). Every figure is generated by a script that also writes its caption and an audit TSV of every plotted value.

## Results

### Direct poly(A) evidence: a scarce, specific channel and one pre-registered operating point

PeakATail calls poly(A) sites (PAS) from the one element of a droplet 3′-end library that observes cleavage directly: the non-templated poly(A) tail carried in a read’s 3′-side terminal soft clip (Fig 1). A read is accepted when it is mapped on the strand of the pass, carries a 16-nt cell barcode and aligns over at most the library read length; it enters the poly(A) channel when its 3′-side terminal soft clip is ≥6 nt and ≥80% A (+) / T (−), with a ≥6-nt run flush to the alignment edge (Fig 1a; per-library parameters in Fig S1). In the measured prototype the flush condition alone buys a 92-fold specificity against wrong-end clips: the identical clip test applied at the read’s opposite (wrong) end fires ∼92× less often. This is a control on alignment artefacts, not on genomic A-runs. Support is counted in distinct (cell barcode, UMI) molecules, never in reads. The channel is scarce by construction: genome-wide, 0.573% of accepted cell-barcoded reads carry a qualifying clip (3, 195, 067 of 557, 564, 408 on the PBMC 10k v3 library; an earlier 1.152% figure was a head-sampling artefact of the QC estimator and is reported only as a corrected estimate). The channel is destroyed outright by any pipeline that trims poly(A) before alignment.

Fig 1 shows what the scarcity buys. On PBMC 10k v3 (CellRanger 3.0.0; 17), 402, 860 candidate peaks enter the funnel; the internal-priming filter removes 68, 855 (17.09%); the survivors split into tier 1 (clip-supported; 167, 629) and tier 2 (coverage-only summits; 166, 376); and the pre-registered default keeps tier-1, internal-priming-pass calls with ≥2 distinct clip molecules (46, 544 rows; 46, 524 scored). Atlas-agreement precision at 100 bp, which is agreement with a strand-matched PolyASite 2.0 (13) representative site and not ground truth, tracks evidence type, not peak geometry: tier 2 scores 0.0558 (n 166, 355), tier 1 at ≥1 molecule 0.3520 (n 167, 565, IP arm), and the default 0.7062, against a gene-body-shuffled null of 0.0217 (peak-level QC in Fig S2).

The internal-priming filter is measured on its own output rather than assumed (Fig 1c). On the PBMC ≥2-molecule arm it removes 15, 588 of 62, 132 calls (25.1%), moving atlas-agreement P@100 from 0.5907 (no-IP arm) to 0.7062 while costing detected-gene recall (R_det) from 0.1911 to 0.1754; the removed calls’ own atlas-agreement precision is 0.246, and 99.97% of removals are triggered by the ≥6-consecutive-A rule. The default is one point on a measured curve (Fig 2a; the full sweep below), and only the ≥2-molecule point was pre-registered.

**Fig 2.**
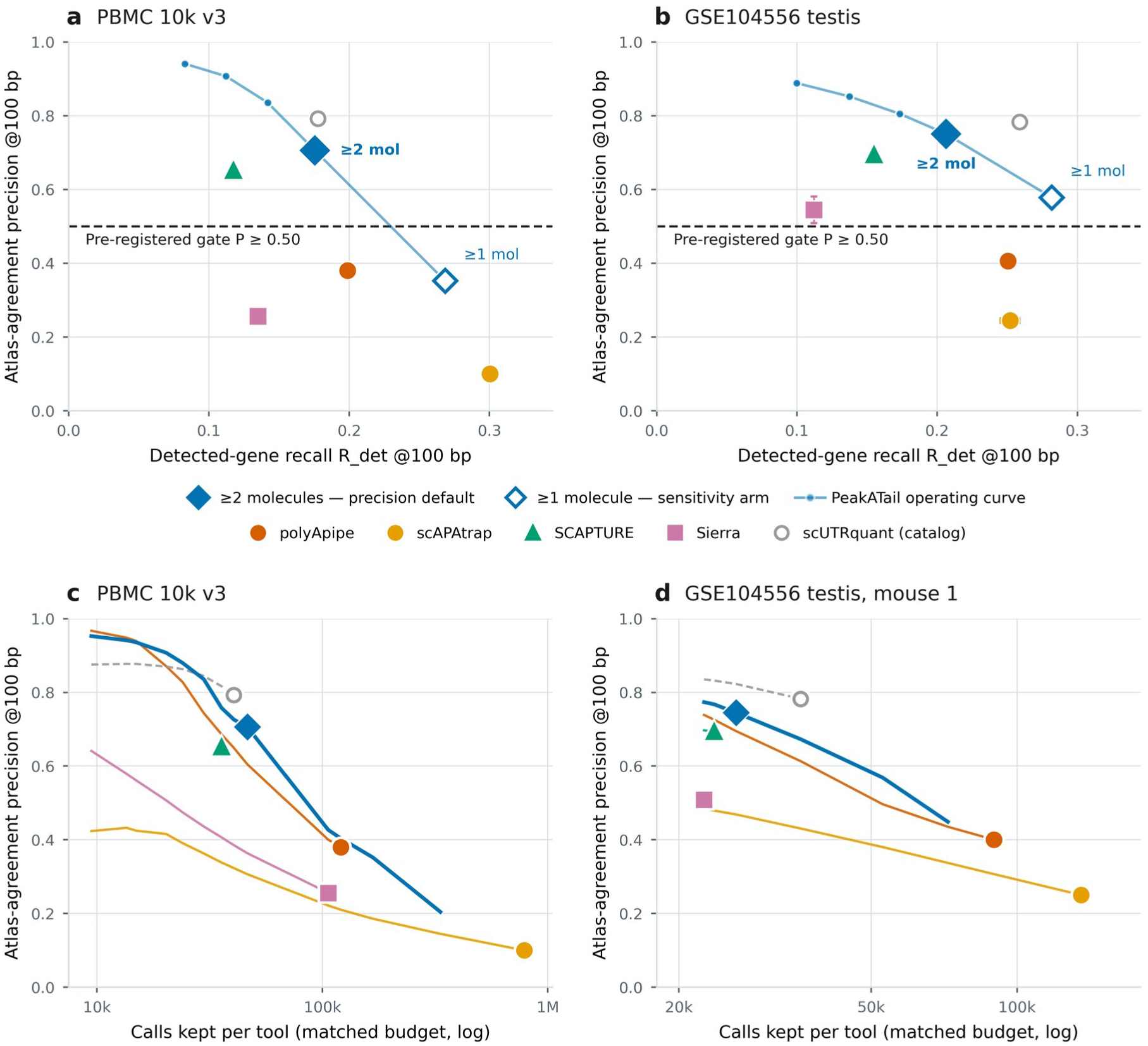
Where PeakATail sits against the field, and what it costs at a matched call budget. One scorer and identical denominators for every tool; 3-seed gene-body-shuffled nulls; PeakATail code 9dfdefb. **(a)** PBMC 10k v3: atlas-agreement precision at 100 bp against detected-gene recall (R_det) at 100 bp. Competitors sit at their own operating points; the light line is PeakATail’s molecule-support operating curve on the internal-priming-filtered arms, where ≥1 / ≥2 / ≥3 / ≥5 / ≥10 distinct clip molecules give precision 0.352 / 0.706 / 0.836 / 0.907 / 0.941 at R_det 0.268 / 0.175 / 0.142 / 0.112 / 0.083. The filled diamond is the pre-registered precision default (tier 1 ∩ internal-priming-pass ∩ ≥2 clip molecules), the open diamond the ≥1-molecule sensitivity arm, and the dashed line the pre-registered gate P@100 ≥ 0.50 (committed 01:19, arms 02:59, v2 re-run 16:14, all 2026-08-21): PASS on all three call sets, 0.7062 (PBMC) and 0.7450 / 0.7572 (mice). Only the ≥2-molecule point was pre-registered; ≥3, ≥5 and ≥10 are descriptive and must never be read as recommendations. **(b)** GSE104556 testis, the same plane, with competitor markers and the two named points as the mean of the two mice with range bars and the curve as the two-mouse mean (precision rising from 0.578 to 0.889 as R_det falls from 0.282 to 0.100). **(c, d)** The same tools at a matched call budget on PBMC 10k v3 **(c)** and testis mouse 1 **(d)**: each ranked call set is truncated to a common number of calls and re-scored, so precision is read at equal N rather than at each tool’s own N; the filled marker on each line is that tool’s own shipped budget (PeakATail 46,524 human / 26,255 mouse 1; polyApipe 120,916 / 89,527; scAPAtrap 787,138 / 135,538). Definitions: atlas-agreement precision (P@100) is the fraction of called 1-bp points with a strand-matched PolyASite 2.0 representative site within 100 bp, i.e. agreement with a curated atlas, not ground truth; R_det denominators are 285,136 detected-gene atlas sites (human) and 126,686 (mouse), and full-atlas recall of the default is 0.089 (PBMC) and 0.098 / 0.100 (mice). Caveats that travel: a single PBMC donor and BAM, and two mice from one study and one chemistry; at the plotted default PeakATail’s recall is below polyApipe’s (−12% / −18%) and its F1_det lead is small (+0.020 / +0.014) at 2.6× fewer calls, so never quote either alone, because at matched call count (panels c, d) PeakATail leads polyApipe on recall at every N tested and on precision at every N from 40,519 upward, and at polyApipe’s own N = 120,916 it leads on both axes (0.4036 / 0.2336 vs 0.3800 / 0.1988); the matched-count precision half is not established for 20,320 ≤ N ≤ 35,759 (competitor-side ties) and is false below N ≈ 15,000, where polyApipe is higher, while the recall half holds at every N; the claim holds among de novo tools only, and catalog-based scUTRquant is drawn as a hollow marker on a dashed line, shown and not ranked; scTail is absent (not runnable on this BAM, read 1 = 28 bp); SCAPTURE’s mouse-2 site-level run completed and scored P@100 0.672, only its per-cell quantification step failed, and that run is never described as invalid; the v1 to v2 −1.1 pp PBMC move is the corrected minus-strand internal-priming window, a correctness change. The atlas-independent corroboration of this same default is Fig S5 (0.7647 of its 46,524 PBMC sites lie within 25 bp of a poly(A)-verified Kinnex long-read 3′ end supported by ≥5 records, against a gene-body-shuffled null of 0.0072); that truth comes from donors other than the short-read library, and its support counts are alignment records that were never de-duplicated. Alt text: Four scatter panels comparing tools on precision against sensitivity. Panels a and b place atlas-agreement precision at 100 bases on the vertical axis against detected-gene recall on the horizontal axis, for PeakATail and five competing tools, on the human PBMC library and on mouse testis respectively; PeakATail’s marker sits highest on the precision axis and to the left of the most sensitive competitor. Panels c and d redraw the same tools at a matched call budget, each tool’s ranked call set becoming a curve through that plane rather than a single point, so the comparison no longer depends on how many calls each tool emits by default.

### Accuracy against the field, under a gate committed before the run

Every comparison in Fig 2 uses one scorer and identical denominators for every tool. Atlas-agreement precision (P@100) is the fraction of called 1-bp points with a strand-matched PolyASite 2.0 representative site within 100 bp (atlases of 569, 005 sites on GRCh38 and 301, 006 on GRCm38); an atlas-novel true site counts as a false positive. Detected-gene recall (R_det) is the fraction of the atlas restricted to genes detected in each dataset that is recovered within 100 bp; the denominators are 285, 136 sites (PBMC 10k v3) and 126, 686 (GSE104556 testis; STARsolo (18, 19); datasets, Fig S1). The precision-first default and its gate, P@100 ≥ 0.50, were committed before the benchmark arms ran (2026-08-21: gate committed 01:19, arms 02:59, v2 re-run 16:14) and passed on all three: 0.7062 (PBMC, n 46, 524), 0.7450 (mouse 1, n 26, 255) and 0.7572 (mouse 2, n 26, 526), each 33–55× its three-seed gene-body-shuffled null (null design, Fig S3); the same gate, unchanged, later passed on a fourth, pre-registered library (Fig S4; below). The −1.1 pp PBMC move from the v1 code is the corrected minus-strand internal-priming window, reconstructed key-for-key.

Across the de novo field, comprising polyApipe (9), Sierra (7), scAPAtrap (8) and SCAPTURE (10), with catalog-based scUTRquant (12) shown but not ranked, call-set sizes span 35, 759 to 787, 138 (22×), and both precision and recall move with call budget, so a single-point comparison measures the operating point, not the tool. (scTail (11) was not runnable on this BAM, whose R1 is 28 bp; SCAPTURE’s mouse-2 run was scored at site level, P@100 0.672.) At its pre-registered operating point PeakATail is the most atlas-concordant de novo call set in the panel on both datasets (atlas-agreement P@100 0.7062 PBMC; 0.7450 / 0.7572 mice). Its recall at that point (R_det 0.1754 / 0.2048 / 0.2080) is below polyApipe’s, but that comparison sets 46, 524 of our calls against 120, 916 of polyApipe’s: at matched call count PeakATail leads polyApipe on recall at every call budget tested and on precision at every budget from 40, 519 calls upward, on both datasets (at polyApipe’s own N of 120, 916, 0.4036 / 0.2336 against 0.3800 / 0.1988), and on both mice the ≥1-molecule arm exceeds polyApipe on precision, recall and F1 simultaneously (0.5686 / 0.2806 / 0.3758 and 0.5880 / 0.2826 / 0.3818 against 0.4005 / 0.2499 / 0.3078 and 0.4119 / 0.2508 / 0.3118). The default is a deliberately conservative point on a curve that dominates polyApipe’s operating point (higher on both axes at the same call count), not the tool’s frontier. Sierra and scAPAtrap were truncated under their own shipped rankings by the identical procedure and sit below PeakATail’s matched-N curve on both axes at every budget tested, on both datasets (scAPAtrap’s rows below its 19, 410-call score cap cut inside tied groups and are not a real ranking, and PeakATail at scAPAtrap’s shipped N is not reachable, since 787, 138 exceeds the caller’s largest possible output of 333, 920). Four bounds travel with this claim: it is restricted to de novo tools (catalog-based scUTRquant is above PeakATail on both axes at several matched budgets, and dropping the restriction makes the claim false); the precision half carries its lower bound (below N ≈ 15, 000 polyApipe’s atlas-agreement precision is the higher of the two), while the recall half holds at every N; SCAPTURE ships no non-circular ranking column, so it cannot be truncated to a matched N; and the precision half is tie-established only where the truncation was re-scored under both within-tie orderings, which was done on PeakATail’s own side at four PBMC and three mouse-1 budgets and not on the competitors’, so between 20, 320 and 35, 759 calls, where the competitor truncations cut inside tied groups, the numbers favour PeakATail but the interval is reported as unestablished rather than as a lead.

The default is also corroborated by truth that owes nothing to the atlas: 0.7647 of its 46, 524 PBMC sites (35, 575; Wilson 95% CI 0.7608–0.7685) lie within 25 bp, strand-matched, of a poly(A)-verified Kinnex (16) x3p long-read 3′ end supported by ≥5 long-read records at the terminus, against a gene-body-shuffled null of 0.0072 (10 seeds; 106.5×); polyApipe’s concordance on the same truth is 0.3895, and the atlas-known, hexamer-pass complement scores 0.8941, calibrating the ceiling of the metric (the pre-registered attempt to certify atlas-novel sites on this truth is the negative result of Fig S5). Two caveats travel with every Kinnex number: the long-read truth is from different donors than the short-read library (donor-mismatched), and its support threshold counts alignment records that were never de-duplicated to molecules, so support thresholds are slightly optimistic. Combining the two truths into one hard false-positive definition: 15.10% of the default’s PBMC calls are false by both truths (no atlas site within 100 bp and no long-read 3′ end at ≥5 records within 25 bp), against 49.50% for polyApipe, on 46, 524 versus 120, 916 calls, so the call-budget caveat applies here as everywhere. The default’s residual error is not internal priming; it is low-support intronic peaks without a poly(A) signal, 35.5% of all its false positives falling in that single class.

Fig S2a decomposes where the default’s stringency acts: 72.2% of tier-1 sites on the PBMC IP arm are single-molecule (121, 041 of 167, 565; 72.1% on the no-IP arm; mice 50.3% / 48.8%), so the ≥2-molecule rule does most of the selection, and the operating curve of Fig 2a, b prices what it costs.

### The trade surface: resolution, reproducibility across libraries, and compute

Sweeping the molecule-support threshold on the IP-filtered v2 arms from ≥1 to ≥10 moves atlas-agreement P@100 from 0.352 to 0.941 on PBMC while R_det falls from 0.268 to 0.083; the mice reach 0.885 / 0.892 at ≥10 (Fig 2a, b). Only the ≥2-molecule point was pre-registered; the others are descriptive, and ≥5 or ≥10 must not be read as a recommendation. The ≥2-molecule default is not the F1 optimum on any dataset: the ≥1-molecule arm reaches F1_det 0.3046 (PBMC) and 0.3758 / 0.3818 (mice) against the default’s 0.2811 and 0.3213 / 0.3264. The default is chosen for reliability, not for F1: it is the arm whose calls a biologist can act on without re-validating each site, and we report both arms as labelled operating points.

Tightening the matching window to 10 bp does not change the de novo atlas-agreement precision ordering (Fig 3a; the recall half of the same window sweep is Fig 3b). The default retains P@10 / P@100 = 0.74 and polyApipe 0.77, against 0.46 (SCAPTURE), 0.33 (Sierra) and 0.24 (scAPAtrap): the atlas-agreement precision lead is not an artefact of a loose window; catalog-based scUTRquant (retention 0.72) sits above the default at every window, again shown, not ranked.

**Fig 3.**
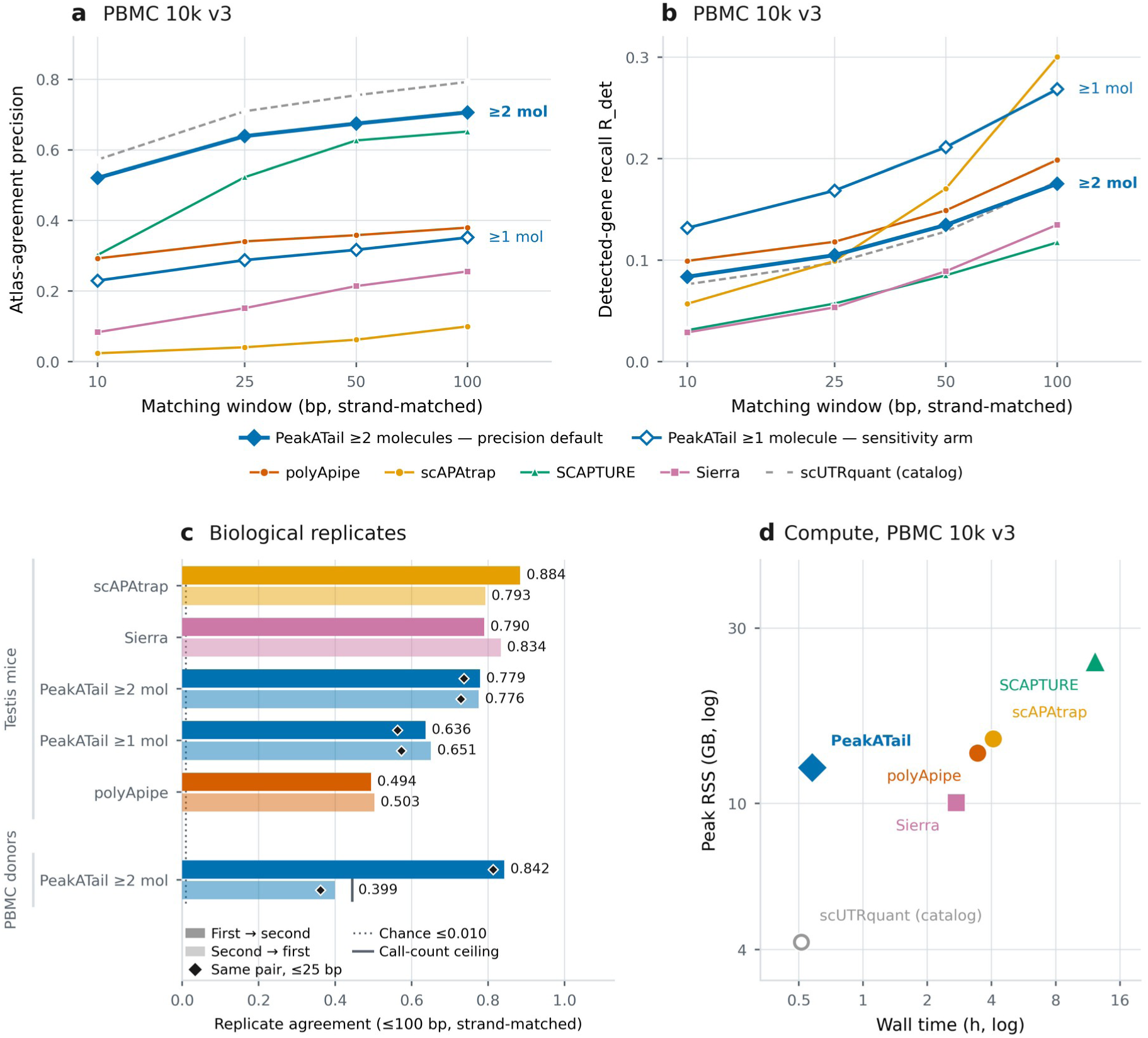
Robustness: matching-window resolution, replicate reproducibility and compute. **(a, b)** Matching-window sensitivity on PBMC 10k v3: atlas-agreement precision **(a)** and detected-gene recall R_det **(b)** at 10 / 25 / 50 / 100 bp for the ≥2-molecule precision default, the ≥1-molecule sensitivity arm and the five competitors. The de novo precision ordering does not change when the window is tightened to 10 bp: P@10 / P@100 is 0.74 for the default and 0.77 for polyApipe against 0.46 (SCAPTURE), 0.33 (Sierra) and 0.24 (scAPAtrap), so the precision lead is not an artefact of a loose window; catalog-based scUTRquant (retention 0.72) sits above the default at every window and is shown, not ranked. **(c)** Replicate agreement: the fraction of one replicate’s call set with a strand-matched call in the other within 100 bp, both directions, with the ≤25 bp value as a diamond. Biological replicates (top block), the two GSE104556 testis mice: default 0.776–0.779 at 100 bp and 0.729–0.737 at 25 bp against the ≥1-molecule arm’s 0.636–0.651, so the ≥2-molecule threshold buys 14.3 / 12.5 points of replicate agreement. Human donors (bottom block), the pre-registered second-donor comparison: 84.2% of donor-2 default calls reproduce within 100 bp in donor 1 (81.3% at 25 bp); the reverse direction, 39.9%, is capped by call-count arithmetic and not by disagreement, because donor 1 emits 46,524 default calls to donor 2’s 20,672, a ceiling of 0.4443 marked by the ceiling tick. **(d)** Compute on the PBMC 10k v3 BAM, log–log: the current caller (34:37, 12.53 GB) beside the five competitors’ own verified runs on the same machine. Caveats that travel: chance agreement in panel c is ≤0.010, and per-tool chance levels span 0.0017– 0.0094 because a denser call set has a higher chance of a nearby match, so the raw ranking is the conservative reading. Honest scope for panel c: the v2 default is above polyApipe and above PeakATail’s own shipped caller but below Sierra (0.790 / 0.834) and scAPAtrap (0.884 / 0.793, a set its own reducePeaks depth-cleaning has already filtered); competitor rows are v1-era measurements against a v2 PeakATail arm; no competitor was run on the second donor, so the human block holds PeakATail arms only and the full pre-registered second-donor protocol is Fig S4; PBMC 10k v3 has no human biological replicate in this benchmark. In panel d the plotted PeakATail point is the clip-seeded arm run *without* the internal-priming filter, the like-for-like pair with v1’s 3:45:53 / 293.7 GB, i.e. 23.5× less peak memory, so no ≥300 GB node is needed; its wall time was measured with all four benchmark arms running concurrently (uncontended single run 27:43): peak resident set size is the production number and no wall time is quoted without its concurrency disclosed. Each competitor row carries its own retry or skipped-stage caveat (Fig S6). The internal-priming-filtered default arm, whose call sets panels a–c show, ran cheaper still (32:59 / 11.12 GB) and is an audit row, not a plotted point. Alt text: Four panels on robustness. Panels a and b are line plots against matching-window width at 10, 25, 50 and 100 bases, showing precision in panel a and detected-gene recall in panel b, with one line for each of three PeakATail operating points and lines for the competing tools. Panel c shows replicate agreement, the fraction of one replicate’s calls with a strand-matched call in the other within 100 bases. Panel d is a log-log plot of run time against peak memory for the current caller beside five competitors.

The two testis mice serve as biological replicates (Fig 3c): 0.776–0.779 of one mouse’s default calls have a strand-matched call in the other within 100 bp (0.729–0.737 at 25 bp), against a chance level

≤0.010, and the ≥2-molecule threshold buys 14.3 / 12.5 points of replicate agreement over the ≥1-molecule arm (0.636–0.651). The honest scope: the v2 default is above polyApipe and PeakATail’s own shipped caller but below Sierra (0.790 / 0.834) and scAPAtrap (0.884 / 0.793, a set its own reducePeaks depth-cleaning step has already filtered), and the competitor rows are v1-era measurements. PBMC is a single donor and a single BAM, so the human replicate had to be created.

We therefore pre-registered a second-library generalisation test before any number from it existed (Fig S4): the frozen v2 tree, run unchanged on pbmc4k (a 10x 3′ v2-chemistry, CellRanger 2.1.0, 98-bp library), with 107 of 108 recorded parameters identical and only the read-length descriptor moving from 91 to 98. The unchanged gate passed at atlas-agreement P@100 0.8279 (n 20, 672), and now holds on four independent libraries (0.7062 / 0.8279 / 0.7450 / 0.7572). One caveat travels with that number: precision generalises, it does not improve. The fixed rule simply lands at a more conservative point on the shallower v2 library: donor 1’s default, cut to donor 2’s call count by any route (its own clip-molecule ranking under both tie-breaks, or a ≥4/≥5-molecule threshold), scores 0.886–0.907 and is above donor 2 on both axes. Donor 2’s R_det is also lower (0.1104 vs 0.1754; denominators 268, 097 and 285, 136 sites; the gap survives every denominator choice). Cross-donor concordance is the human analogue of Fig 3c: 84.2% of donor 2’s default calls are reproduced in donor 1 within 100 bp (∼361× the genic-shuffle null); the reverse direction is 39.9% against its 44.4% arithmetic ceiling, because donor 1 emits 46, 524 default calls to donor 2’s 20, 672; both directions are reported, neither alone. 10x publishes no donor identifier for pbmc4k: this is demonstrably a second library, chemistry and CellRanger version, only presumptively a second individual.

The same code revision that produced the v2 numbers removed the tool’s hardware barrier (Fig 3d; detail in Fig S6). On the PBMC BAM, peak RSS fell from 293.7 GB to 12.53 GB (23.5×), so the ≥300 GB node requirement is gone, and wall time fell from 3:45:53 to 34:37 on the like-for-like pair (the clip-seeded arm without the internal-priming filter); the v2 wall time was measured with all four benchmark arms running concurrently (∼28–35 min; uncontended single run, 27:43). At cohort scale (17 libraries from 14 patients, ∼224 GB of BAM) the identical configuration fell from 9:06:26 to 1:06:26 (8.2×; v2 run with --peak-workers 16) at identical output, 505, 197 unified PAS from both codes, with peak RSS falling from 13.63 to 13.53 GB.

### The recall budget, measured

We measured how much of the recall gap any configuration of this caller could ever recover. Of the 235, 111 detected-gene atlas sites the PBMC default misses, 79.96% have no peak of any kind within 100 bp (84.53% in mouse 1), so at most 20.04% / 15.47% of the gap is reachable by any threshold, filter or tier policy; keeping every peak the caller ever produced reaches R_det 0.3406 (PBMC) and 0.3278 (mouse 1), the hard ceiling for anything downstream of this caller’s peak calling. That ceiling is a property of PeakATail’s evidence-generation (seeding) step, not of the library: what the reads could support for a different candidate generator is not bounded by it. The denominator itself deserves context rather than being read as fully attainable: only 44.6% of the detected-gene atlas is corroborated by matched-tissue Kinnex long reads at ≥5 records (the union count, 127, 188 of 285, 136). Those long reads come from a different donor, with un-deduplicated record support, so "not corroborated" is not "not real", and the fraction contextualises the denominator without bounding what is real. Against the x3p-corroborated subset (107, 260 sites) the same default recalls 0.4372, not 0.1754. Those are facts about two different subsets and must not be divided across each other. The atlas places 20.0 sites in an average detected human gene where the default emits 4.0: absolute recall against such a denominator should not be read as sensitivity.

### The switch test is calibrated because the shipped defaults were not

Before reporting a single switch we asked whether `peakatail switch diff` controls the false-discovery rate it states (Fig 4). A label-permutation harness on testis mouse 1 (1, 230 cells with expression-derived stage labels, orthogonal to the PAS matrix under test; 76, 711 PAS; three stage pairs) ran 20 permutations, shared identically across configurations, through the full pipeline (six configurations × 21 runs, zero failures): Fisher on read counts, Fisher on cell counts and a pairwise negative-binomial test, each with v2’s shipped top-200 Wilcoxon marker pre-selection on and off.

**Fig 4.**
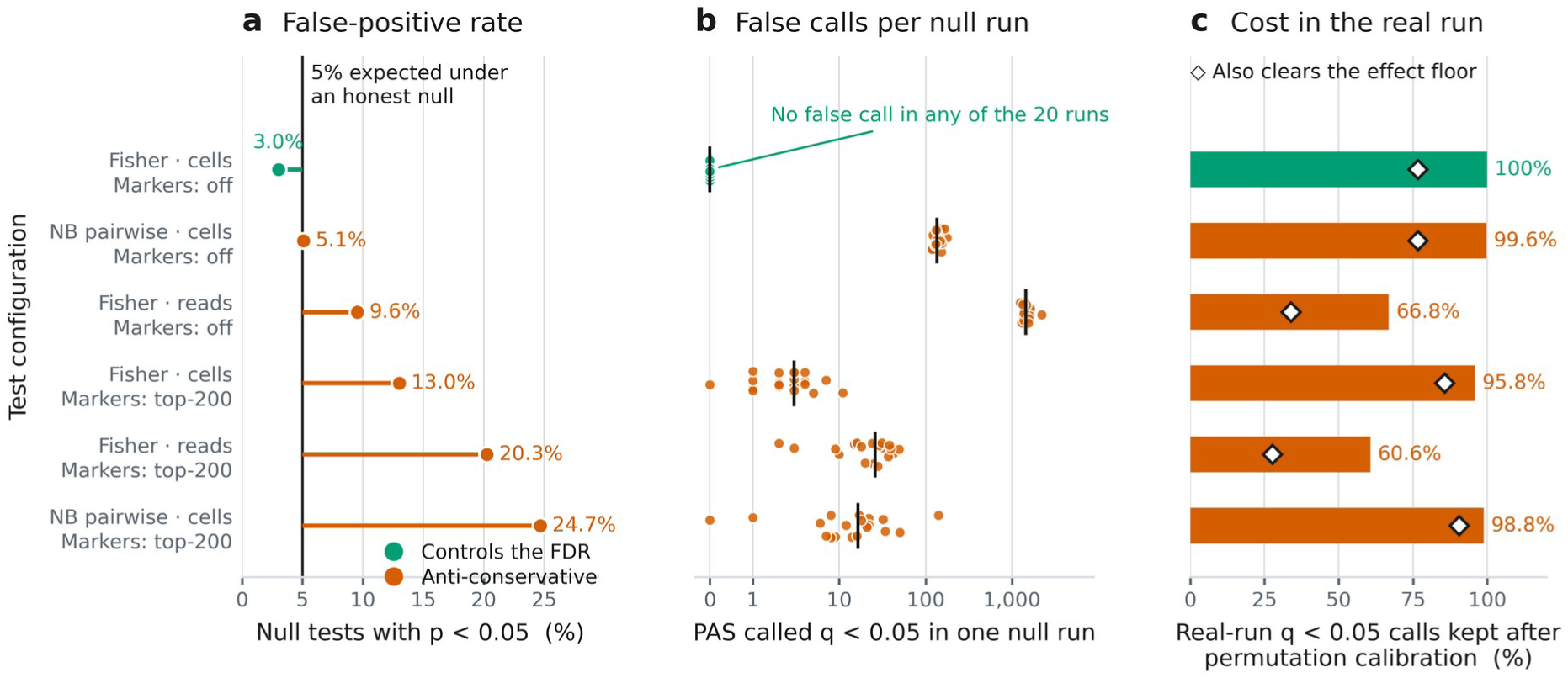
Which switch-test configurations control the false-discovery rate under label permutation. Each row is one configuration a user chooses on the command line: statistical test × count unit (`--count-mode reads|cells`) × marker pre-selection (`--marker-top-n 200`, the default shipped at v2 `9dfdefb` / version 0.2.0, or `0`; that default has since been flipped to `0` upstream, version-bumped to 0.3.0). All six are scored against the same label-permutation null run through the identical pipeline: testis mouse 1, 1,230 cells with expression-derived stage labels orthogonal to the PAS matrix under test (SPC 370 / RS 504 / ES 356), 76,711 PAS, three stage pairs, 20 permutations shared identically across configurations, the full `peakatail switch diff` pipeline re-run per permutation. The score is therefore a property of the configuration, not of one dataset. **(a)** The share of null tests reaching p < 0.05 in each configuration against the 5% an honest null produces (vertical rule); the stem runs from that reference to the observed rate, so its length and side are the size and the sign of the miscalibration. **(b)** False calls made by a *single* null run: one dot per label permutation (20 per configuration), bar = median, symmetric-log axis so exact zeros are drawn at 0. **(c)** The cost in the real, unpermuted comparison: the share of each configuration’s nominal q < 0.05 calls that survive permutation-calibrated q (bar; label = surviving count / all real-run calls), with a diamond marking the share that also clears the pre-registered effect floor (|Δproportion| ≥ 0.1 for Fisher, |log2FC| ≥ 1 for the negative binomial). Result: every arm that keeps the top-200 marker pre-selection that v2 shipped as its default is anti-conservative (null p < 0.05 of 20.3% for Fisher on reads, 13.0% for Fisher on cells and 24.7% for the negative binomial, with 19–20 of 20 null runs carrying a q < 0.05 call), as are Fisher on reads (9.6%) and the negative binomial (5.1%) without it; Fisher on cells with pre-selection off is the one arm that controls the false-discovery rate (3.0% null p < 0.05, 0.00% null q < 0.05, 0 of 20 null runs with any hit), conservative rather than exact. Caveats that travel: every switch result here is at `--isoform-agg per_gene`, so a "switch" is a shift in a PAS’s share of its gene and not necessarily tandem 3′UTR polyadenylation, and the two are not the same event; re-running the same cells, labels and test at `--isoform-agg within_utr`, which scopes the denominator to the PAS sharing a 3′UTR isoform, reproduces 90.9% and 91.6% of the per-gene calls in the two mice (96.8–97.1% of within-UTR calls are also per-gene calls; 161,874 and 165,964 PAS × stage-pair tested by both), so the scope changes under a tenth of the calls but does change which event is being named; that comparison required repairing the released caller, in which `--isoform-agg within_utr` aborted on every invocation (two defects on a path with no test coverage, both repaired upstream after the freeze together with the sibling `between_utr` output), so no PeakATail version published at the time of the analysis could run it, and the first release that can is 0.3.0; marker pre-selection also computes the within-gene Fisher denominator from the marker-restricted matrix, so marker-on and marker-off are different tests rather than subsets, and their real-run hit counts are not comparable; one mouse and one tissue with very large true stage effects, so real-run hit counts are detectability upper bounds, not precision; the label-permutation null tests exchangeability under the global null only; a Kolmogorov–Smirnov test against uniform rejects for every arm because Fisher p-values are discrete (mass at p = 1), and is reported, not gated; "calibrated" is our pre-stated operational rule (null p < 0.05 ≤ 7%, p < 0.01 ≤ 1.5%, q < 0.05 ≤ 5%, ≤ 25% of null runs with any hit), not a community standard. Per-configuration diagnostic depth is Fig S7. Alt text: Three panels, each row within a panel being one switch-test configuration the user selects on the command line. Panel a, horizontal bars giving the share of label-permuted tests reaching a nominal p below 0.05 in each configuration, against a vertical reference line at the 5 percent an honest null would produce; most configurations extend well past the line and one falls below it. Panel b, a dot plot with one dot per label permutation and a median bar per configuration, showing the false calls produced by a single null run. Panel c, the share of each configuration’s nominally significant calls in the real, unpermuted comparison that survive the stricter treatment.

The shipped defaults are anti-conservative: with pre-selection on, 20.3% (Fisher/reads), 13.0% (Fisher/cells) and 24.7% (negative binomial) of label-shuffled tests reached p < 0.05, and 19–20 of 20 null runs carried a q < 0.05 call (Fig 4a, b). Exactly one configuration controls the FDR: Fisher on cells with pre-selection off, at 3.0% null p < 0.05, 0.00% null q < 0.05 and no q < 0.05 call in any of 20 null runs. It is conservative rather than exactly calibrated, and never anti-conservative in any stage pair or expression stratum (2.94–3.13% and 2.6–4.0%; Fig S7a–b).

The mechanisms are named and separately demonstrated. Marker pre-selection is a label double-dip: markers are chosen on the labels under test, and restricting the calibrated configuration’s null p-values to PAS selected on permuted labels reproduces the inflation (17.4%). It also computes the within-gene Fisher denominator from the marker-restricted matrix, making marker mode a different test, not a subset. Fisher on read counts pseudoreplicates UMIs within cells, inflated even without pre-selection (9.6% null p < 0.05, ∼1, 500 false q < 0.05 calls per null run). The negative-binomial test is tail-inflated by its plug-in dispersion floor: in the pre-selection-off arm, tests at the 10⁻⁴ floor are 8.5% of its null tests yet carry 67% of its null q < 0.05 hits (13% in the marker arm; Fig S7e).

PeakATail therefore changed its shipped configuration to Fisher on cells with pre-selection off ( `--marker-top-n 0`, nominal Benjamini–Hochberg q; 20) as its only default, and requires permutation-calibrated q whenever pre-selection or the negative-binomial test is used (Fig S7d). That flip is post-freeze and does not describe the code behind this paper: at v2 ( `9dfdefb3`) `--marker-top-n` still defaults to 200, and the default was changed to 0 only after the freeze. A flagless `ema switch diff` at this paper’s own frozen commit (`peakatail switch diff` from 0.3.0) therefore runs the anti-conservative 13.0% arm, and every switch result reported here required passing `--marker-top-n 0` explicitly; `--strategy fisher` and `--count-mode cells` are already the defaults at `9dfdefb`, so the marker flag is the only one a reproducer must add. "Calibrated" is our pre-stated operational rule (null p < 0.05 ≤ 7%, p < 0.01 ≤ 1.5%, q < 0.05 ≤ 5%, ≤ 25% of null runs with any hit), not a community standard, and the assessment rests on one mouse and one tissue with very large true stage effects. The harness measures only this tool’s test: no competitor’s differential test has been run through it, so Background’s field-level concern about uncalibrated shipped defaults is, for every other tool, a hypothesis this paper does not test.

### A biological control with its own nulls: the spermatogenesis gradient

We next asked whether the calibrated test recovers a known biological program when every claim carries its own null: both testis mice, the frozen merged caller, expression-derived stage labels, spermatogonia excluded in advance as fragile (Fig 5). The claim, in the exact form that survives adversarial verification on GSE104556 (21, 22):

> Per-gene 3′UTR usage shifts progressively proximal from spermatocyte to round spermatid to elongating spermatid in both mice: 31.4% and 30.5% of depth-guarded genes shorten monotonically across the three stages against label-shuffle nulls of 17.4% and 21.0% (z = 10.5 and 6.4, 20 shuffles), exceeding monotone lengthening (486 vs 318 and 366 vs 291 genes), and the composition-controlled per-cell distal-usage residual falls at every step (Cliff’s δ SPC vs ES = 0.55 and 0.63, outside the entire label-shuffle null range). The gradient is a per-gene, equal-weight statement: the per-gene medians are not monotone (RS marginally above SPC) and the across-gene UMI-weighted distal index reverses between RS and ES, where protamine transcripts alone carry ∼25% of the UMIs at ceiling PDUI.

**Fig 5.**
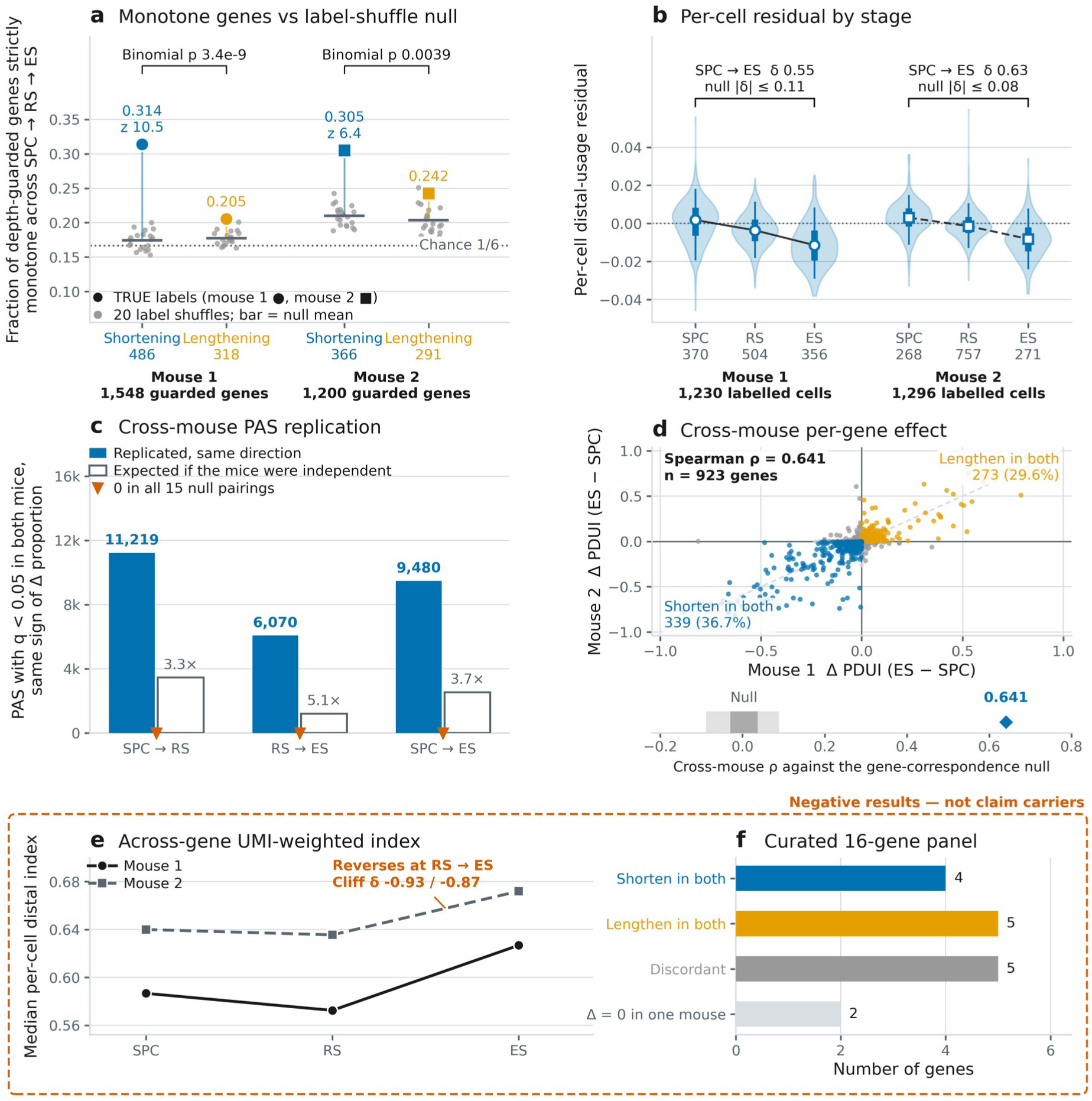
Per-gene 3′UTR usage shifts progressively proximal from spermatocyte (SPC) to round spermatid (RS) to elongating spermatid (ES), in both mice. GSE104556 testis, both mice, frozen merged caller 9dfdefb3; stage labels are marker-panel argmax over Leiden clusters of the STARsolo gene-expression matrix, orthogonal to the PAS matrix under test. PDUI = distal / (proximal + distal), higher = longer 3′UTR; the statistic is the depth-weighted pseudobulk per-gene PDUI over genes with ≥50 UMI at the PAS pair in every stage (1,548 / 1,200 genes). **(a)** The claim carrier: the fraction of depth-guarded genes strictly monotone across the three stages, per mouse and per direction, with the 20 label-shuffle draws as grey dots (bar = null mean) and the true value as a filled marker: shortening 0.314 vs 0.174 ± 0.013 (z = 10.5) and 0.305 vs 0.210 ± 0.015 (z = 6.4); lengthening 0.205 vs 0.177 ± 0.010 (z = 2.8) and 0.242 vs 0.204 ± 0.019 (z = 2.1). The dotted line is 1/6, the chance rate for a random ordering of three stages; the bracket carries the direction-specific evidence, the excess of shortening over lengthening (486 vs 318, binomial p 3.4e-9; 366 vs 291, p 0.0039). **(b)** The composition-controlled per-cell index, for each cell the mean over genes of (per-cell PDUI − that gene’s three-stage pseudobulk PDUI), falls at every step in both mice: Cliff’s δ SPC vs RS 0.26 / 0.32, RS vs ES 0.41 / 0.46, SPC vs ES 0.55 / 0.63, against 20-shuffle null ranges of [−0.10, +0.11] and [−0.08, +0.07]. Violin = all cells, thick bar = interquartile range, whisker = 5th–95th percentile, open marker = median. **(c)** PAS called switching at q < 0.05 in both mice with the same sign of Δproportion (Fisher, cells mode, no marker pre-selection), per stage pair: 11,219 / 6,070 / 9,480 of 36,840 / 35,668 / 35,382 coordinate-matched PAS, sign agreement 99.7–99.8%, 3.25–5.05× the independence expectation (hollow bars), and 0 replicated in all 15 null pairings; the calibration re-validates on this caller (pooled null p < 0.05 3.169% / 3.156%, 0 q < 0.05 hits in all 30 null Benjamini–Hochberg families). **(d)** The per-gene effect ΔPDUI (ES − SPC) in mouse 1 against mouse 2 over the 923 genes guarded in both: Spearman ρ = 0.641 (Pearson r = 0.842) against a gene-correspondence null of 0.004 ± 0.033 (200 permutations, max |ρ| 0.088; strip below the scatter), with 339 genes (36.7%) shortening in both mice, 273 (29.6%) lengthening in both, 253 strictly discordant and 58 with Δ exactly 0 in one mouse. **(e, f)** Boxed negative results, not claim carriers. **(e)** The across-gene UMI-weighted per-cell distal index *rises* between RS and ES in both mice (0.587, 0.572 and 0.627 across SPC, RS and ES in one mouse, and 0.640, 0.636 and 0.672 in the other; Cliff’s δ RS vs ES −0.93 / −0.87), because protamine transcripts alone carry 25.2% (mouse 1) and 14.5% (mouse 2) of the guarded ES UMIs at ceiling PDUI. **(f)** Of the 16 genes of the curated literature panel measurable in both mice, only 4 shorten in both (Prm3, Ybx2, Ppp1cc, Nsun7), 5 lengthen in both (Prm1, Tnp2, Odf1, Spata19, Smcp), 5 are discordant and 2 uninformative. Caveats that travel: monotone lengthening is also above its null, because a label shuffle destroys ordered structure of either sign, so never write "shortening is above null" without the lengthening column, and quote the excess; the per-gene medians are not monotone (RS sits marginally above SPC in both mice) and must never be quoted as the gradient, which is carried by panel a, the paired per-gene tests and the per-cell residual of panel b; the statistic is the depth-weighted pseudobulk PDUI and that choice must be stated wherever these numbers are used, the cell-weighted arm being unusable at this depth; a bare Wilcoxon p from this design is not interpretable without its shuffle-null column; spermatogonia are excluded from the claim, because they lengthen under exactly this labelling on 64 / 68 cells; gene-level and across-gene UMI-weighted summaries disagree in direction (panels a–d against panel e) and both are reported, and across-gene UMI weighting is not used for the claim; per-gene reproducibility and net shortening are different claims and must not be conflated (ρ 0.641 against 339 both-shorten vs 273 both-lengthen, binomial p 0.0085); replication here is two mice of one study, one protocol and one chemistry with argmax expression labels, so animal-level noise is controlled and protocol-level artefacts are not; the switch test’s null is conservative rather than exact, so the true hit counts are not a count of true switches, and cross-mouse PAS matching is coordinate-based, so only the matched subset (∼60% of tested PAS) can replicate; the ≥50-UMI depth guard biases the surviving gene set toward highly expressed genes (the shuffle null uses the same guarded set); the literature panel is a named-gene measurement and runs through the PAS-to-gene assignment that was defective in overlapping loci at the frozen commit (repaired upstream after the freeze, with a documented residual, and none of that code is behind this figure), which is not excluded for individual panel verdicts; and the 3′UTR-scoped PDUI arm could not be produced by the frozen code and is absent here (a defect repaired upstream after the freeze; that fix is not in 9dfdefb3 and so is not in the code behind this figure). Alt text: Six panels covering both mice. Panel a, the fraction of depth-guarded genes whose 3’UTR usage is strictly monotone across the three sperm-development stages, per mouse. Panel b, a composition-controlled per-cell index by stage. Panel c, sites called as switching in both mice with the same direction of effect. Panel d, a scatter of the per-gene effect in mouse 1 against mouse 2 over the genes guarded in both. Panels e and f are boxed negative results rather than claim carriers: panel e an across-gene per-cell distal index, panel f the small number of genes from the curated literature panel that shorten in both mice.

Three cautions are bound to the claim. Monotone lengthening is also above its null (z = 2.8 and 2.1), because a label shuffle destroys ordered structure of either sign, so the direction-specific evidence is the excess of shortening over lengthening: strong in mouse 1 (binomial p 3.4e-9), clear in mouse 2 (p 0.0039; 0 of 40 null draws, one excess draw per shuffle per mouse with both mice’s 20 shuffles pooled, reached the excess in absolute value). The z-scores are read against a 20-shuffle null distribution, whose empirical tail resolves only to p ≤ 1/21; beyond that floor the tail claim rests on the z statistic itself (a 200-permutation null for this arm is a named pending item, not part of this run’s record). The ≥50-UMI depth guard keeps 1, 548 and 1, 200 genes; the per-cell residual Cliff’s δ is 0.554 and 0.632 exactly (Fig 5a–b).

Fig 5c, d carry the reliability payload. The calibrated test re-validates on this caller (pooled null p < 0.05 3.169% / 3.156%; 0 q < 0.05 hits in all 30 null BH families and 0/10 null runs), and its calls replicate across animals: 6, 070 / 9, 480 / 11, 219 PAS switch at q < 0.05 in both mice in the same direction for the ES–RS / ES–SPC / RS–SPC pairs (sign agreement 99.7–99.8%), zero replicated PAS in all 15 null pairings, and per-gene effects correlate across mice (ΔPDUI ES−SPC Spearman ρ = 0.641, n = 923 genes guarded in both; gene-correspondence null 0.004 ± 0.033).

Two results ran against expectation; Fig 5e, f box them as findings, not footnotes. The across-gene UMI-weighted distal index reverses between RS and ES in both mice (Cliff’s δ −0.93 / −0.87): protamine transcripts at ceiling PDUI alone carry 25.2% and 14.5% of guarded elongating-spermatid UMIs, so gene-level and UMI-weighted summaries disagree in direction and both are reported. And of a 27-gene panel of spermatid-expressed and 3′-end-processing genes assembled for this control, 16 were measurable in both mice: only 4 shorten in both, 5 lengthen in both, 5 are discordant and 2 are uninformative. The panel does not reproduce under our PAS-to-gene assignment, is dropped as a positive control, and only the per-gene, null-referenced statement stands. The panel is our own assembly and not a published gene list, so its failure is not a failure to reproduce anyone’s reported panel; what it tests against is the general finding of widespread 3′UTR shortening across mouse spermatogenesis (5). One alternative explanation is not excluded here: a named-gene measurement runs through the PAS-to-gene assignment that was defective in overlapping loci in the code that produced these numbers (a defect known at the time of the freeze and repaired only afterwards; nothing here is recomputed under the repair), so per-gene mis-assignment could contribute to individual panel verdicts. Replication here is two mice of one study and one chemistry: it controls animal-level noise, not protocol-level artefacts.

### Reliable cell-type APA switches in a tumour cohort

We then applied the same discipline where labels are noisier and effects smaller (Fig 6). The Laughney lung-adenocarcinoma cohort (23, 24) comprises 17 libraries from 14 patients; the replication primary is 15 libraries from 12 patients (the pre-registered MetBone exclusion; one further library has a single confirmed cell type, so no testable pair). The unit of replication is the patient, never the library, so two-library patients cannot "replicate" within one person.

> Across 12 patients (15 scRNA-seq libraries) of the Laughney lung-adenocarcinoma cohort, processed in a single cohort run so that all samples share one PAS identifier space, PeakATail called cell-type APA switches in 59 cell-type pairs over a pre-registered precision-first PAS universe (clip-supported, internal-priming-filtered, ≥2 clip molecules; 80, 464 sites). Requiring a switch to be called (Fisher on cells, BH q<0.05) in the same direction in at least two independent patients, with any opposite-direction patient vetoing the call, 15, 942 of 2, 128, 711 tested (pair, PAS) hypotheses replicate (0.75%), of which 15, 212 also exceed an effect floor of |Δproportion| ≥ 0.1; these are 5, 951 distinct PAS in 2, 883 genes across 47 pairs, and 10, 415 (cell-type pair, gene) combinations. Under ten patient-wise label-shuffle nulls run through the identical pipeline, no feature replicated in any combination. Requiring three patients retains 5, 438 PAS-level switches.

**Fig 6.**
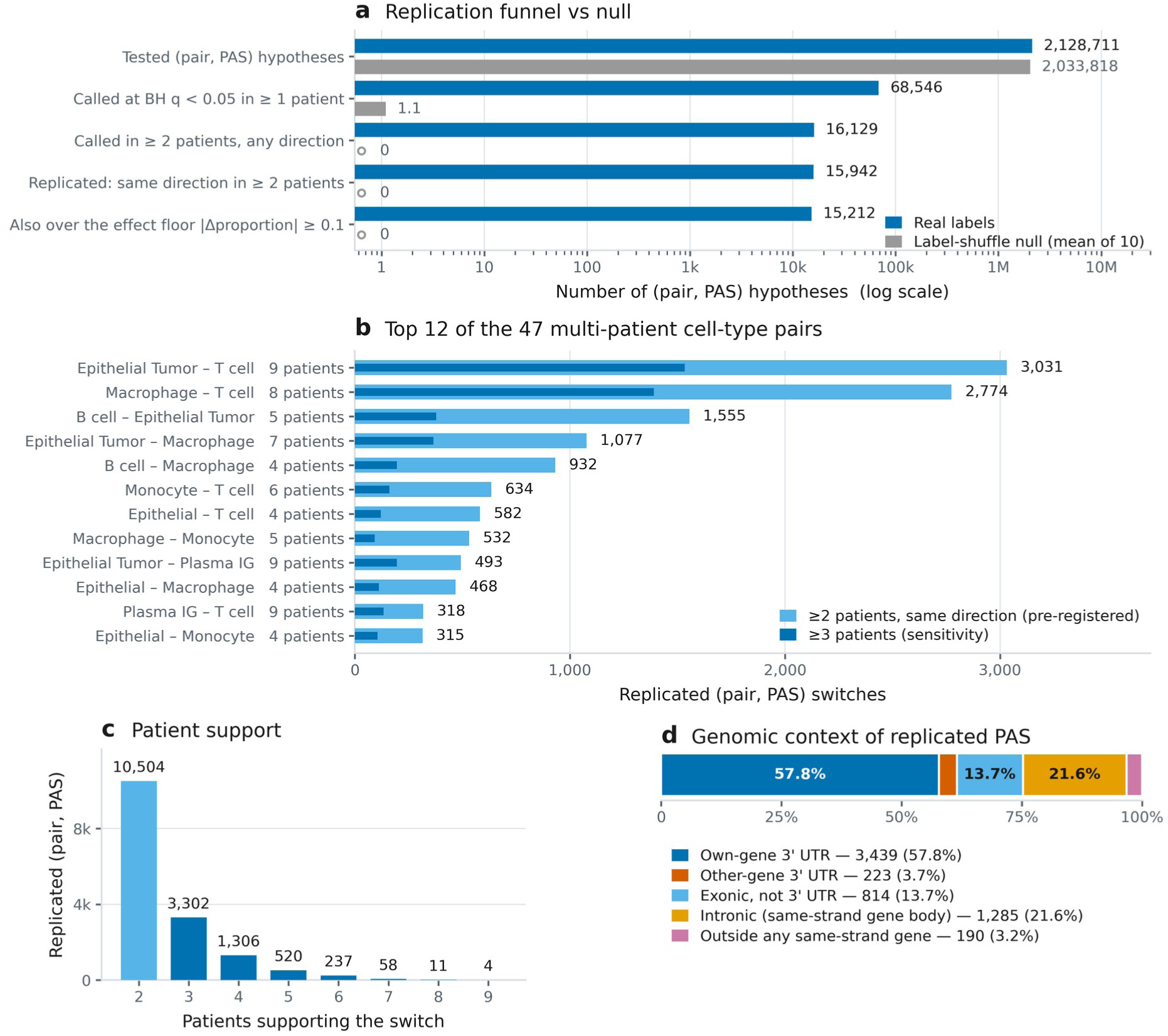
Reliable cell-type APA switches in a tumour cohort. Fifteen scRNA-seq libraries from 12 patients of the Laughney lung-adenocarcinoma cohort, peak-called in a single run together with the other two libraries of the 17-library cohort, so that all libraries share one PAS identifier space. Cell counts are cohort-wide where the curation was done: 29,063 curated cells over all 17 libraries, 18,651 label-confirmed (64.2%) under the pre-registered policy, of which the 17,373 falling in the 15 primary libraries entered the tests; 59 cell-type pairs over a pre-registered precision-first PAS universe (clip-supported, internal-priming-filtered, ≥2 clip molecules; 80,464 sites); test = Fisher on cells with no marker pre-selection; the unit of replication is the patient, so the three two-library patients count once each. **(a)** The replication funnel on one log axis, real (blue) against the mean of ten patient-wise label-shuffle nulls (grey): 2,128,711 (cell-type pair, PAS) hypotheses were tested. Of these, 68,546 were called at Benjamini–Hochberg q < 0.05 in at least one patient, 16,129 were called in at least two patients in any direction, and 15,942 replicated in the same direction with no opposite-direction patient; of the 187 lost at that step, 148 reached no single direction in two patients and 39 were vetoed by a patient going the other way. A further restriction to calls over the |Δproportion| ≥ 0.1 floor leaves 15,212. The null tests a comparable 2,033,818 hypotheses per combination and reaches a mean of 1.1 single-patient calls (range 0–4) and zero at every later stage. **(b)** The top 12 of the 47 cell-type pairs tested in ≥2 patients, at the two-patient threshold (light) with the three-patient subset overlaid (dark); "n pt" is the number of patients in which the pair was testable. The other 35 multi-patient pairs hold 3,231 more switches (1–296 each), and 12 pairs were tested in a single patient and can never replicate. **(c)** Patient support: the three-patient sensitivity set is exactly the ≥3-patient tail (5,438), and 10,504 of the 15,942 replicated switches (66%) sit at the two-patient minimum. **(d)** Honesty panel, the genomic context of the 5,951 distinct replicated PAS, drawn as an exclusive strand-matched partition: 57.8% in the assigned gene’s own 3′UTR, 3.7% in another gene’s 3′UTR, 13.7% exonic but not 3′UTR, 21.6% intronic and 3.2% outside any same-strand gene, i.e. 96.8% inside a same-strand gene body and 61.5% in some same-strand 3′UTR. Caveats that travel: the two 3′UTR figures must never be conflated, and the own-gene figure (57.8%) is the one to use whenever the claim is about gene-level interpretation; that fraction, like the 2,883-gene and 10,415 (pair, gene) counts, is computed through the PAS-to-gene assignment that was defective at `9dfdefb` (repaired upstream after these results were frozen, so the counts reported here retain the defect and would change on a re-run), while every PAS-keyed statistic is unaffected; the null resolves only to an empirical p ≤ 0.091, the ten-permutation floor, so report "none in ten label-shuffle nulls" and never an FDR; no ranked list of named top-switch genes may be published from these runs, because 12 of the top-30 gene-level rows name a gene whose 3′UTR does not contain the representative PAS; the assignment fix landed upstream after this paper’s v2 freeze at `9dfdefb`, so it is in none of the code that produced any number here, and such a list becomes publishable only once the cohort is re-run on the fixed assignment — and then only with the recorded residual, that `in_utr` still tests a gene-length footprint anchored at the gene terminus rather than real UTR intervals, so a minority of contested assignments (96 of 465 relabels in the fix’s own audit) still resolve by terminus proximity; the pre-registered effect floor is applied per patient and the patients are then re-counted, so it removes 730 switches, from 15,942 to 15,212, more than the 291 whose consensus effect alone falls below 0.1, so never quote the consensus-below-floor count as the floor’s yield; the MetBone exclusion is pre-registered but its stated premise was wrong (the near-zero clip rate behind it was a head-of-BAM sampling artefact), and the 13-patient sensitivity run differs by 0.4%, confined to one pair; pairs tested in more patients replicate more, so panel b tracks cohort composition as much as biology, and two thirds of the replicated set rests on the two-patient minimum (panel c), the weakest support the pre-registration allows; one library trips the per-run null rule only through Benjamini–Hochberg discreteness (3 hits in 9.1 M null tests) and is kept and disclosed; one cohort, one chemistry, one caller, with no orthogonal 3′-end assay confirming these sites; these are v2-code numbers, and the v1-code chain recorded 14,480 replicated over a 74,954-PAS universe and remains reproducible as a labelled comparison. Alt text: Four panels on the tumour cohort. Panel a, the replication funnel drawn on a single logarithmic axis, the real counts beside the mean of ten patient-wise label-shuffle nulls, the null bars falling to zero where the real bars do not. Panel b, horizontal bars for the top 12 of the 47 cell-type pairs tested in at least two patients, shown at the two-patient threshold with the three-patient threshold overlaid. Panel c, the distribution of patient support across replicated sites. Panel d, an honesty panel giving the genomic context of the distinct replicated sites as an exclusive strand-matched partition.

The two test totals connect as follows: the switch test itself runs per library, and the ten patient-wise label-shuffle combinations re-run those per-library tests in full (58.9 M library-level null tests across the 15 primary libraries), while the 2, 128, 711 hypotheses above are the distinct (cell-type pair, PAS) rows entering the patient-level replication filter. The 58.9 M null tests yielded 11 nominal q < 0.05 calls, none co-occurring twice: not within a combination, not across combinations, not in two libraries of one patient. We report this as none in 10 label-shuffle nulls (empirical p ≤ 0.091, the 10-permutation floor); it is a null control, not an FDR estimate. Per-library null calibration is conservative (pooled null p < 0.05 0.84–2.73%); one library, GSM3516672-StageIB, trips the pre-registered per-run count rule (3/10 permutations with a q < 0.05 hit vs an allowed 2, from BH discreteness in the largest library; all four rate criteria pass), disclosed because "all libraries pass calibration" would be false.

Two disclosures bound Fig 6. The MetBone exclusion stands because it was pre-registered, but its stated premise was wrong: the near-zero clip-rate estimate behind it came from a QC estimator sampling only the first 200, 000 cell-barcoded reads of a coordinate-sorted BAM. MetBone in fact carries 306, 202 clip molecules (30.2% of its 28, 983 post-tier-filter PAS hold ≥2 of its own clip molecules); the sensitivity run including it differs by 0.42%, confined to the one pair it contributes. Second, no ranked list of named top-switch genes is reported: in overlapping loci, 12 of the top-30 gene-level rows (40%) have a representative PAS outside the named gene’s 3′UTR and 8 of 30 (27%) sit in a different gene’s 3′UTR, so switch results stay keyed by PAS identifier, with replication statistics unaffected; the overlapping-loci half of that issue was reworked upstream after this paper’s freeze (the repair resolves PAS-to-gene ties from the annotation instead of terminus proximity and recovers CD68 on chr17), it is not in the v2 code that produced these numbers, and its authors disclose a residual — the 3’UTR test still uses a fixed-length footprint anchored at the gene terminus, so 96 of 465 chr17 relabels are still decided by proximity — so the gene list stays withheld here and its release would require both that assignment and a re-run of these results on it. Fig 6d gives the genomic context of the 5, 951 replicated PAS: 96.8% inside a same-strand gene body, 61.5% in some same-strand 3′UTR, 57.8% in the assigned gene’s own 3′UTR. The gene-keyed quantities, namely 2, 883 genes, the 10, 415 (pair, gene) combinations and the 57.8% own-3′UTR fraction, are computed through that same assignment and carry the same caution as the withheld gene list; the gene-body and any-same-strand-3′UTR fractions and every PAS-level statistic do not depend on choosing an assigned gene.

### What did not work, reported as results

We pre-registered four claims and their acceptance gates before the final run existed and report all four outcomes (Figs S5 and S8). Three negative results are contributions in their own right; each names the alternative explanation tested and excluded.

### The pre-registered "trusted de novo PAS" definition failed its gate (Fig S5)

The definition was clip-supported, ≥2 molecules, internal-priming-filtered, canonical hexamer at −40..−5 and ≥100 bp from any PolyASite 2.0 (13) site (the pre-registration also named PolyA_DB, hg19-only, which could not be used); it was required to reach ≥70% concordance with poly(A)-verified Kinnex long-read 3′ ends within 25 bp (16), at ≥5 supporting long-read records at the terminus (site-level truth from different donors). Its 7, 259 sites (15.6% of the PBMC precision default) reached 48.5% (95% CI 47.4–49.7). The failure is a property of the definition: re-running it unchanged after the two caller fixes moved it down from 52.1% to 48.5%; every alternative truth set misses at the pre-registered window (GEM-X 51.5%, pooled 58.2%); and donor mismatch is excluded, since atlas-known hexamer-pass sites score 89.4% on the same truth. The loss is localised to the atlas-novelty stage, which falls from 0.811 to 0.485 ( the hexamer adds nothing among atlas-novel sites, hexamer-fail 0.487): 69.4% of trusted-novel sites are intronic, and 27.0% sit within 25 bp of a Kinnex internal-priming decoy terminus against 7.6% for atlas-known, hexamer-pass sites. A stricter genomic A-richness rule does not fix this: added on top of the shipped filter, the Kinnex-style rule removes 31 of 46, 524 default sites, a no-op. The residual class therefore survives both rules, and the diagnosis, chosen after seeing the data, remains a hypothesis. The sites are 65–179× enriched over a gene-body-shuffled null, but the target was an absolute fraction, and no site in this paper is called "trusted novel".

### The clustering-novelty claim was withdrawn by its own ablation (Fig S8)

Cells cluster on PAS-site profiles into expression-derived cell types (PBMC AMI 0.708, replicated in 17/17 cohort libraries at median AMI 0.662), but collapsing the 275, 370 sites to 14, 891 per-gene totals recovers the same types (AMI 0.698). Site-level resolution adds nothing, so the signal is 3′-end expression re-encoded rather than isoform choice; clustering ships as an optional convenience, and no claim rests on it.

### Pre-registration, reported with its failures

Every acceptance gate was committed before the run it judges, with git-verified commit times where they exist. The record covers every gate and every arm, reports both failures, discloses as post hoc the one sweep that was post hoc, and marks the single still-untested pre-registration pending rather than omitting it.

### Curated-atlas benchmarking rewards atlas-shaped priors

In algorithmic-headroom work, a per-site scoring model trained against the curated atlas won 15.7 points of atlas-agreement precision over the shipped ≥2-molecule rule at matched call count (P@100 0.8634 against 0.7062) while losing 7.5–10.8% of long-read-verified sites at every operating point held equal, and reached the rule’s truth:decoy odds only at 84, 737 calls, where its long-read precision is 17.7 points below the rule’s (Supplementary Table T8). A transferable variant likewise fooled the atlas while losing 15% of long-read-verified sites. A model can raise the benchmark number while destroying agreement with evidence the atlas never saw. Two rules follow, in force throughout this paper: long-read truth is a read-out only, never used to select a threshold, feature or model; and every later parameter pre-registration carries an anti-atlas-prior guard: the long-read truth must move in the same direction before a change ships as a default. No learned score ships here; the reported default remains the pre-registered rule, and its call files are byte-for-byte identical across the caller versions tested.

## Discussion

What a precision-first default buys a biologist is trust per call. At its pre-registered operating point (clip-supported tier-1 sites, internal-priming-filtered, ≥2 distinct clip molecules), PeakATail’s output is the most atlas-concordant de novo call set in the benchmark on both datasets and is corroborated by donor-mismatched long reads (Fig 2); the matched-call-count analysis, with the N-range bounds that travel with it, is given once in Results and is the only licensed form of the head-to-head. The two-truth hard-false-positive composite of Results is the summary a biologist should carry: 15.10% of the default’s PBMC calls fail both truths, against 49.50% of polyApipe’s (9) at 2.6× the call count. The default is not the F1 optimum on any dataset (Fig 3): it is chosen for reliability, being the arm whose calls a biologist can act on without re-validating each site, and both arms ship as labelled operating points.

The cost is recall, and both halves of the trade run into measured ceilings (Results, "The recall budget, measured"): the poly(A)-clip evidence channel is 0.573% of accepted cell-barcoded reads (correcting our earlier documented 1.152%); roughly four fifths of the detected-gene atlas sites the default misses have no peak of any kind within 100 bp, capping what any threshold or tier policy downstream of peak calling could recover; and under half of the detected-gene atlas is itself corroborated by donor-mismatched long reads. Absolute recall against a pan-tissue atlas should not be read as sensitivity.

The negative results carry the field-level lessons. First, curated-atlas benchmarking rewards atlas-shaped priors: the atlas-trained per-site scoring model of Results improved against the truth set while degrading against the biology, so every gate here pairs atlas agreement with an atlas-independent long-read check, and the field’s benchmarks should too. Second, pre-registration repeatedly caught our own overclaims: the sweep that informed the ≥2-molecule threshold is disclosed as post hoc; created after a stale run failed the original gate, it informed a choice and is cited nowhere as a result; and the adoption criterion for the revised defaults returned FAIL at Δ = 0.000000 because it was drafted against a default configuration that never existed; the outputs were byte-identical, an error only a pre-written criterion could expose. The same discipline produced the paper’s other negatives: only one of six switch-test configurations controls the false-discovery rate, and the pre-registered trusted-novel definition reached 48.5% long-read concordance against its 70% target, reported as a failure (Fig S5).

Limitations. The human results rest on one donor at depth; the second library (pbmc4k), whose pre-registered default gate returned PASS at atlas-agreement P@100 0.8279 with cross-donor site concordance of 84.2% within 100 bp (descriptive, not gated; Fig S4), carries no published donor identifier and is presumptively, not provably, a second individual. The two testis mice come from one study and one chemistry. The Kinnex truth is donor-mismatched and site-level: it corroborates positions, not donor-specific usage. Two defects were open when these results were produced and are present in them; both were closed upstream only after the frozen commits, so neither fix is in this paper’s numbers. PAS-to-gene assignment in overlapping loci was wrong in v2 (repaired only after the freeze, and the maintainer records the repair as still approximating 3′UTR footprints by gene-terminus distance in a minority of ties,), so cohort switches here are keyed by PAS identifier and no ranked gene list appears here; and the per_isoform switch mode had a degenerate-pair defect in v2 (repaired upstream after the freeze) and underlies no result here. A third cited defect is still open at the time of writing, the switch test’s default marker pre-selection: upstream has since flipped the shipped default to `--marker-top-n 0` and taken the Fisher denominator from the full count matrix, while a label-independent pre-filter, the `nb_pairwise` dispersion floor, the `within_utr`/`between_utr` denominator, two documentation defects and the run-manifest commit hash remain. It bears on no result reported here, every one of which was run with marker pre-selection off.

Future work follows the measured roadmap. Cross-sample "cohort borrowing" is the one recall mechanism with verified headroom: mouse-1 single-clip-molecule tier-1 sites corroborated by a mouse-2 default call within 10 bp score atlas-agreement P@100 0.6683 versus 0.4144 for expression-matched singletons. But internal priming replicates across samples too, the effect is unmeasured on human data, and it ships only behind its own pre-registration. The 3′UTR singleton promotion is pre-registered (rule, thresholds and falsifiers fixed, flag default OFF) and untested: its deciding evidence is reserved to data unused in its discovery, and its ceiling is ∼+0.04 R_det. Re-ranking is nearly exhausted: a long-read-trained score gains +14.3% relative R_det at matched atlas-agreement precision on the one library it was fitted on, but adopting it would replace the pre-registered default and so requires its own pre-registration and multi-dataset runs; the next real gain lies in the candidate generator or in the chemistry, not in thresholds.

PeakATail calls poly(A) sites from direct poly(A)-tail read evidence and reports, for every claim, the gate it was required to pass, the null it was tested against, and whether it failed. Under that discipline its precision-first default is the most atlas-concordant de novo call set in the benchmark on both datasets, corroborated by donor-independent long reads, and its calibrated, replication-filtered switch test yields cell-type APA switches that recur across patients (replication primary: 15 libraries from 12 patients) with none in 10 label-shuffle nulls (empirical p ≤ 0.091, the 10-permutation floor). Just as deliberately, it reports a failed pre-registered gate, a withdrawn clustering claim, anti-conservative shipped defaults and a spermatogenesis gene panel that does not reproduce. We offer the method, and the evaluation discipline it was built to survive, as the contribution.

## Data availability

### Tool code

PeakATail is developed openly at https://github.com/BMGLab/PeakATail and archived on Zenodo under the concept DOI 10.5281/zenodo.22697820, which always resolves to the latest release; the released versions carry their own DOIs (0.3.0, 10.5281/zenodo.22697821; 0.3.1, 10.5281/zenodo.22699241). No released version corresponds to the code that produced the results reported here, which predates the 0.3.0 release and is identified by the two frozen commits named below; the release DOIs are given so that the software itself is citable, not as a substitute for those commits. The frozen commits used in this paper are `4efeb125` (v1) and `9dfdefb3eb353b0817ef79c4eb9ace6d6c8aab53` (v2); both are public in that repository.

### Analysis code and manuscript record

The benchmark scorer (`score_tool.py`), the replication filter (`replication_filter.py` v0.2.0), the long-read truth-set build scripts, the label-confirmation and universe policy files, every figure-generating script with its audit TSVs and script-written captions, the pre-registration documents with their appended amendments, and the run manifests are in the companion repository https://github.com/BMGLab/Project_PeakATail. The repository is public and MIT-licensed. The derived Kinnex truth and decoy point sets, which have no upstream accession of their own, are distributed in it.

### Data

All sequencing data are previously published or publicly distributed: the 10x Genomics public libraries `pbmc_10k_v3` and `pbmc4k` (BAMs from the 10x public data portal), the mouse testis libraries of GEO accession GSE104556, and the 17 lung-adenocarcinoma libraries of GEO accession GSE123904. The accuracy reference is the PolyASite 2.0 atlas (GRCh38 and GRCm38 representative sites). The PacBio Kinnex long-read truth and decoy point BEDs (x3p and GEM-X) are distributed with the companion repository together with the script that builds them.

Not applicable. This study analyses only previously published, publicly available datasets; no new human or animal data were collected.

## Supplementary data

Supplementary Data are available at *NAR Genomics and Bioinformatics* Online.

Full supplementary figure legends, with their travelling caveats, are in the Supplementary Data; the titles below fix the frozen numbering. The Supplementary Data also contain a formal specification of the method: every quantity reported here, stated in notation and read off the frozen release, with one table giving the value of every parameter at that release. It restates the Methods below and extends nothing: no quantity it defines underlies a number the Methods does not also describe in prose.

**Fig S1 The datasets behind every headline number, with per-library poly(A) clip rates.** Unmeasured cells are shown as missing, never approximated; head-sample estimator values are comparable only with each other, never with the genome-wide rate; 1.152% appears only as the superseded reference.

**Fig S2 v2 peak-call QC and the tier-2 cleavage-offset census.** Every count names its arm; the ∼90–105 nt tier-2 offset is real, tier-2-specific, and its correction fails every pre-registered acceptance criterion, a negative disposition; the census ran on development slices, so full-BAM generalisation is an inference.

**Fig S3 Why the benchmark is built this way: dump vs curated regimes, chance = reference density, one reconciled null.** The retired-regime values are shown only as the discrepancy being reconciled and may not be quoted as results.

**Fig S4 Second-donor validation (pre-registered): pbmc4k gate PASS at 0.8279, cross-donor concordance both directions.** The matched-N control travels with it: precision generalises, it does not improve; a second library, chemistry and CellRanger version, only presumptively a second individual.

**Fig S5 The pre-registered trusted-novel definition and its failure: 48.5% long-read concordance against a 70% target.** A negative result, written as one; no site in this paper is called "trusted novel"; enrichment over the shuffled null (65–179×) does not rescue an absolute-fraction target; the residual class survives both the shipped and the Kinnex-style internal-priming rule.

**Fig S6 Compute detail behind Fig 3d**. No wall time is quoted without its concurrency disclosed; peak RSS is the production number; competitor retry/skip caveats travel verbatim.

**Fig S7 Calibration deep-dive behind Fig 4**. Per-pair and per-stratum null rates, permutation-calibrated q, reads-vs-cells pseudoreplication, the NB dispersion-floor diagnostic; single mouse, single tissue; the label-permutation null tests exchangeability only.

**Fig S8 The dropped clustering claim, stated honestly.** The positive result never appears without its ablation on the same axis: per-gene totals recover cell types as well as site-level profiles, so no novelty claim rests on clustering.

**Supplementary Table T7 Competitor tool versions, parameters, run commands and run-log caveats.** Every version string is read from the `tool_version.txt` each run wrote at run time, and every invocation is the run script actually executed; both are in the analysis repository at the paths the table gives.

## Acknowledgements

The authors thank Ebru Kocakaya for her work on the benchmarking and analyses reported here, and Melike Güler for useful discussions and for volunteering to perform tests in the initial stages of this project.

## Author contributions

A.A.T. developed the software and designed and implemented the poly(A)-site calling method. E.B.Z. contributed to the software and to testing it. Y.K. conceived and supervised the study, acquired the resources for it, and carried out the benchmarking and the downstream analyses. All authors designed the evaluation protocol and its pre-registered gates, interpreted the results, wrote the manuscript, and read and approved the final version.

